# Basal forebrain activity predicts functional degeneration in the entorhinal cortex and decreases with Alzheimer’s Disease progression

**DOI:** 10.1101/2023.03.28.534523

**Authors:** Marthe Mieling, Martin Göttlich, Mushfa Yousuf, Nico Bunzeck, the Alzheimer’s Disease Neuroimaging Initative

## Abstract

**BACKGROUND AND OBJECTIVES:** Recent models of Alzheimer’s Disease (AD) suggest the nucleus basalis of Meynert (NbM) as the origin of structural degeneration followed by the entorhinal cortex (EC). However, the functional properties of NbM and EC regarding amyloid-β and hyperphosphorylated tau remain unclear.

**METHODS:** We analyzed resting-state (rs)fMRI data with CSF assays from the Alzheimer’s Disease Neuroimaging Initiative (ADNI, n=71) at baseline and two years later.

**RESULTS:** At baseline, local activity, as quantified by fractional amplitude of low-frequency fluctuations (fALFF), differentiated between normal and abnormal CSF groups in the NbM but not EC. Further, NbM activity linearly decreased as a function of CSF ratio, resembling the disease status. Finally, NbM activity predicted the annual percentage signal change in EC, but not the reverse, independent from CSF ratio.

**DISCUSSION:** Our findings give novel insights into the pathogenesis of AD by showing that local activity in NbM is affected by proteinopathology and predicts functional degeneration within the EC.

## Introduction

The basal forebrain’s nucleus basalis of Meynert (NbM) has recently been suggested as the origin of structural degeneration in Alzheimer’s disease (AD) followed by the entorhinal cortex (EC) and other cortical brain regions ^1,2^. For instance, grey matter loss was more prominent in the NbM compared to the EC in cognitively healthy humans with an abnormal CSF biomarker of amyloid-β (Aβ) and hyperphosphorylated Tau (pTau). Moreover, the NbM’s baseline volume predicted the longitudinal structural degeneration in the EC, further suggesting a trans-synaptic spread of Aβ starting in the NbM ^1,2^. This observation in humans is in line with animal work and adds a crucial upstream link to the subsequent spread from EC to other medial temporal lobe structures, including the hippocampus, and more distal neocortical brain regions such as the posterior parietal cortex ^1,3–6^. Importantly, evidence in favor of such a pathological staging model is mainly limited to anatomical studies, and, therefore, the functional properties of both the NbM and EC during the disease progression of AD in humans remain unclear.

Since functional brain changes in AD often precede structural degeneration ^7–9^, we investigated the functional properties of the NbM and EC, including their functional connectivity. To this end, we used data from the Alzheimer’s Disease Neuroimaging Initiative (ADNI) and performed a longitudinal region of interest (ROI) analysis over 2 years, focusing on regional and interregional resting-state functional MRI (rsfMRI) properties. In detail, we analyzed (a) the fractional amplitude of low-frequency fluctuations (fALFF) to quantify spontaneous neuronal activity ^10–12^, (b) regional homogeneity (ReHo) reflecting the synchronicity of neural activity between a voxel and its neighboring voxels ^13^, and finally (c) the functional connectivity between NbM and EC. While all three measures may help to gain new insights into AD progression, we initially focused on fALFF given its established role ^14–16^, and report ReHo and functional connectivity analyses in the supplementary material.

In the first step, baseline signals and longitudinal functional changes were compared based on harmonized CSF assays of Aβ and pTau in NbM and EC. Subsequently, we investigated functional changes in disease progression using the CSF markers. Finally, we tested the competing models NbM◊EC vs. EC◊NbM on a functional level. Our main hypothesis was that functional signals in the NbM predict functional change in EC, which would provide further evidence supporting the pathological staging model originating from NbM to EC. From a more general perspective, we aimed to provide new insights into the underlying functional properties of AD, which may contribute to further developing markers and treatment strategies.

## Methods

### ADNI data

Data used in the preparation of this article were obtained from the Alzheimer’s Disease Neuroimaging Initiative (ADNI) database (adni.loni.usc.edu). ADNI was launched in 2003 as a public-private partnership, led by Principal Investigator Michael W. Weiner, MD, to test whether serial magnetic resonance imaging (MRI), positron emission tomography (PET), other biological markers, and clinical and neuropsychological assessment can be combined to characterize the progression of mild cognitive impairment (MCI) and early Alzheimer’s disease (AD).

Since rsfMRI was not acquired in all ADNI cohorts, here data were combined from ADNI-GO, ADNI-2 (ADNI-GO/2) and ADNI 3, downloaded from the Image and Data Archive (IDA) platform run by the Laboratory of Neuro Imaging (LONI) (https://ida.loni.usc.edu). Specifically, we only selected data from participants with CSF biomarkers, and two rsfMRI scans acquired with a delay of two years with the same MR scanner and head coil to ensure within-subject comparability.

### Image acquisition

Participants were scanned at multiple sites equipped with 3-Tesla MRI scanners according to unified ADNI monitoring protocols ^17^. To ensure maximum compatibility between the measurements, we followed ADNI’s recommendations and included only the basic rsfMRI version but not advanced version of ADNI 3 since it is not compatible with ADNI-GO/2. Moreover, all participants here were examined with the same scanner and head coil for both timepoints, T1 and T2 (https://adni.loni.usc.edu/methods/mri-tool/mri-analysis/). Further, we only included MRI data with excellent, good, or fair quality. For further information on image acquisition, see the supplementary material and http://adni.loni.usc.edu.

### Data preprocessing

Considering their specific scanning parameters such as TR, slice order, and volume number, all data were preprocessed with the Data Processing Assistant for Resting-State fMRI Advanced (DPARSFA, http://rfmri.org/dpabi) toolbox version 5 (release 5.2_210501), which is based on the Statistical Parametric Mapping toolbox (SPM 12, https://www.fil.ion.ucl.ac.uk/spm/) for MATLAB®. It started with the removal of the first ten volumes and subsequently included the following steps a) slice time correction; b) spatial realignment; c) T1 co-registration to the mean functional image; d) CSF, gray and white matter tissue segmentation, and spatial normalization using diffeomorphic anatomical registration using exponential lie algebra (DARTEL) ^18^ for T1 images; e) regression of nuisance variables; f) normalization to MNI space and resampling to an isotropic voxel size of 3 mm of the functional images using the parameters estimated by DARTEL (see supplementary material for a detailed description).

To reduce the influence of excessive head motion, participants exhibiting more than 3.0 mm of maximum movement and a 3.0-degree rotation angle were discarded. Further, images were visually inspected after co-registration, segmentation, and normalization to guarantee high quality. This included a specific focus on signal loss and artifacts in our regions of interest (NbM, EC) by overlaying a ROI mask in standardized space; especially, the EC represents a region that might often be affected by artifacts ^19^. For a detailed description of the preprocessing steps, excluded participants, ROI definition, and rsfMRI analyses for fALFF, ReHo, and the functional connectivity, see supplementary material.

### CSF biomarker

AD neuropathology includes the accumulation of Aβ resulting in plaques and pTau leading to neurofibrillary tangles ^20,21^. To better understand how both relate to functional degeneration in NbM and EC, we followed previous studies ^1,2^ and used ADNI’s CSF samples, produced with a fully automated Elecsys® protocol of Aβ and pTau from the first measurement (T1). For each participant, we extracted Aβ 1-42 and pTau181 values. Since the protocols by Elecsys® are still under development, the results are restricted to a specific technical limit (>1700 pg/mL). Higher values were provided by extrapolation of the calibration curve for research purposes only but not diagnostics. Further information on CSF draws and analyses can be found at http://adni.loni.usc.edu.

Here, we analyzed both proteins by using a previously established ratio of pTau / Aβ, which is known to highly concord with PET measures and clinical diagnoses ^23,25^. Based on these findings, the standardized and cross-validated cut-off of 0.028 was used to divide the participants into an abnormal (pTau / Aβ ≥ 0.028) and a normal (pTau / Aβ < 0.028) CSF group ^1,23,25^. Importantly, no participant classified with AD had a normal CSF ratio, but a few (n=10) participants classified with MCI did, which indicates an unclear etiology. Nevertheless, we included them based on biological instead of a syndromal grouping ^26^.

### Neuropsychological assessment and clinical diagnosis

All participants underwent a comprehensive neuropsychological test battery. Here, T1 scores are used, including validated memory (MEM) and executive function (EF), based on a confirmatory factor analysis ^27,28^. Memory scores include the AD Assessment Scale, Logical Memory test, Mini-Mental-State Examination (MMSE), Rey Auditory Verbal Learning Test (RAVLT). EF scores are based on the Category Fluency, Digit Span Backwards, Digit Symbol Substitution, Trails A and B, and the Clock Drawing tests ^27,28^. We were also interested in the Montreal-Cognitive-Assessment (MoCA), Sum of Boxes in the Clinical Dementia Rating Scale (CDRSB) and the Alzheimer’s Disease Assessment Scale Cognitive (ADAS-Cog 13) to get a deeper understanding of the participants’ cognitive profiles (see below).

Furthermore, we included participants’ T1 diagnosis made by the ADNI Clinical Core: cognitive normal (CN) (CDR=0, MMSE=24-30), mild cognitive impairment (MCI) (CDR=0.5, MMSE=24-30), and Alzheimer’s disease (AD) (CDR=0.5-1, MMSE=20-26). These classifications represent widely used cognitive and functional measures in clinical trials ^29–31^. Further information regarding diagnostic is available at http://adni.loni.usc.edu.

### Participants

We included rsfMRI data from ADNI-GO/2 and ADNI 3 – but, importantly, only those that also offered a subject’s CSF draw (see below) temporally related to a rsfMRI acquisition (e.g., a participant’s screening MRI and baseline lumbar punction measurement). This measurement served as T1 measurement in the analyses. To maximize the number of subjects, the second measurement was selected after an interval of 1.5 years ±12 months (T2) ^1^. Further details on inclusion and exclusion criteria for participating in ADNI are available under http://adni.loni.usc.edu. In total, 153 participants for ADNI-GO/2 and 141 for ADNI 3 (only basic rsfMRI version) fulfilled our inclusion criteria. However, a large proportion had to be excluded mainly based on fMRI data quality (see supplementary material). Thus, data from n=71 participants were analyzed, which could be further subdivided into those with normal CSF (nCSF, n=37) and abnormal CSF (aCSF, n=34) values (Table 1).

**Table 1.**
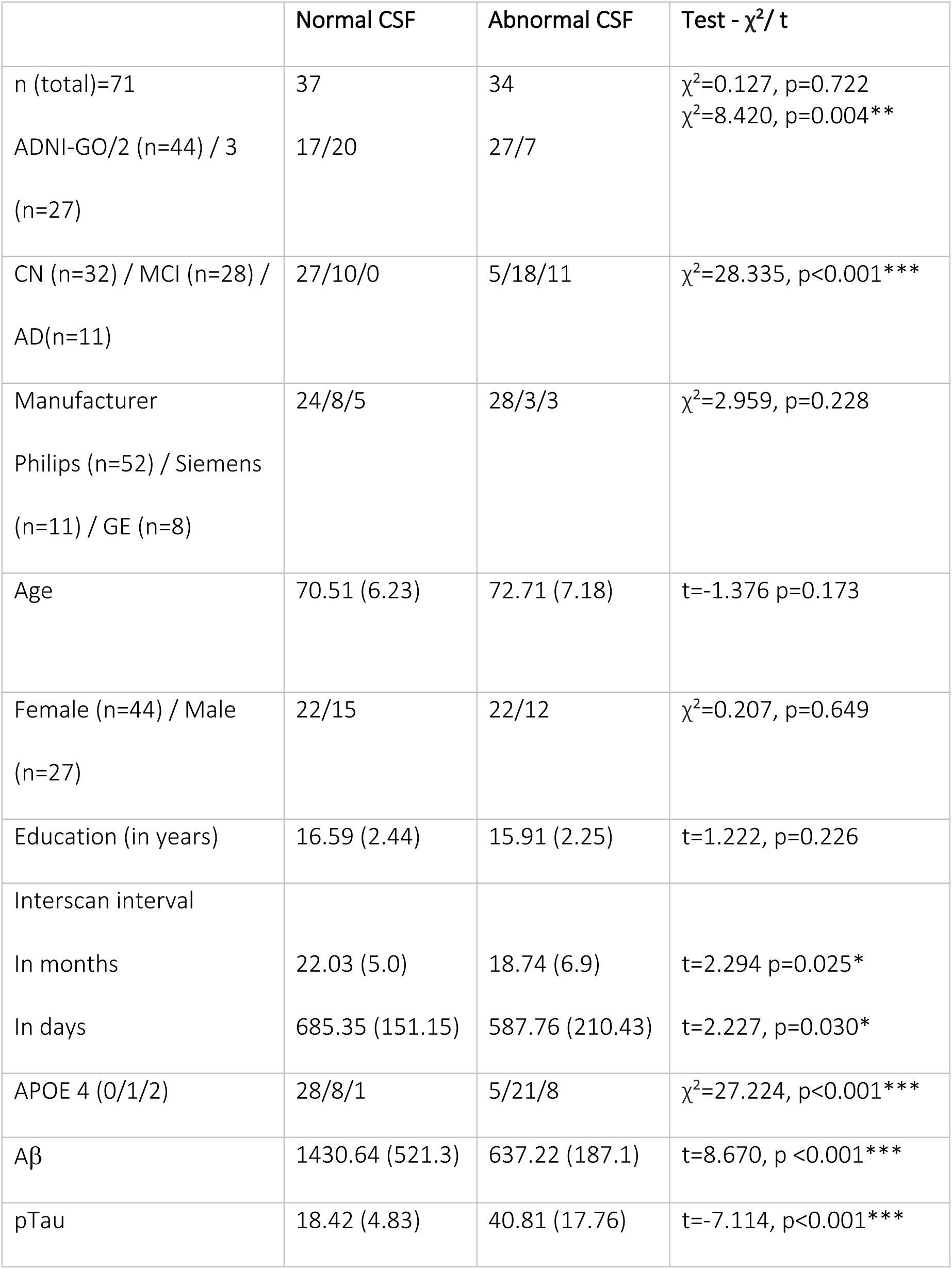
Participants’ demographics and information on APOE4 genotype and harmonized CSF assays. Information of the final sample from ADNI-GO/2 and ADNI-3 grouped by CSF. Means and standard deviation (SD) are represented and the respective t-test or chi-square test to investigate possible group differences. Baseline clinical diagnosis: CN=cognitive normal; MCI=mild cognitive impairment; AD=Alzheimer’s Disease. Age and education were assessed in years. APOE4 status: no allel / 1 allel / 2 allels. Aβ=amyloid-β in pg/ml as concentration of the amyloid-β 1-42 peptide. pTau=in pg/ml as CSF concentration of hyperphosphorylated tau. *p<0.05, **p<0.01, ***p<0.001

Table 1 gives an overview of the participants’ demographics, as well as information on APOE4 genotype and harmonized CSF assay, and Table 2 shows the neuropsychological test results at baseline (T1).

**Table 2.** Neuropsychological test results at baseline, compared by CSF normal vs. abnormal. The mean values with standard deviation (SD) of normal vs. abnormal CSF groups. The abnormal CSF group reflected worse performance in all neuropsychological tests. MEM: memory function score; EF: executive function score; MMSE: Mini-Mental State Examination; ADAS-Cog 13: Alzheimer’s Disease Assessment Scale-Cognition Subscale, 13 tasks; CDRSB: Clinical Dementia Rating Scale; MoCA: Montreal-Cognitive-Assessment; Clock drawing: clock drawing test * significant after Bonferroni correction p < 0.05/n (n = 7 tests).

| Neuropsychological<br>testing | CSF groups (mean (SD)) |  | F- value | P- value |
| --- | --- | --- | --- | --- |
|  | Normal | Abnormal |  |  |
| MEM score | 0.88 (0.6) | -0.08 (0.97) | F(1,66)= 23.4 | <0.001* |
| EF score | 1.02 (0.76) | -0.16 (1.1) | F(1,66)= 23.8 | <0.001* |
| MMSE | 29.08 (1.12) | 25.82 (3.50) | F(1,66)= 23.35 | <0.001* |
| ADAS-Cog 13 | 9.8 (4.8) | 22.89(14.26) | F(1,66)= 26.82 | <0.001* |
| CDRSB | 0.3 (0.55) | 2.63 (2.36) | F(1,66)= 34.72 | <0.001* |
| MoCA | 25.89 (2.34) | 20.91 (5.72) | F(1,66)= 22.5 | <0.001* |
| Clock drawing | 4.76 (0.55) | 4.03 (1.22) | F(1,66)= 8.73 | 0.004* |

### Ethics

Each center collecting data for ADNI provided an IRB (Institutional Review Board) approval and meets ADNI’s requirements. Informed consent was obtained from all ADNI participants (for more information at http://adni.loni.usc.edu). The analyses presented here were approved by the local Ethics Committee of the University of Lübeck and carried out after ADNI’s recommendations including the approval of the manuscript before submitting to a journal.

### Data availability statement

All data are freely available upon request from the Image and Data Archive (IDA) run by the Laboratory of Neuro Imaging (LONI) (https://ida.loni.usc.edu).

### Statistical analyses

#### Mixed ANCOVA

Mixed ANCOVAs were carried out for all measures separately (i.e., fALFF, ReHo) to compare baseline signals and the annual percentage signal change (APSC, see below) between regions (NbM and EC as a within-subject factor) and CSF groups (normal and abnormal as a between-subject factor). Covariates such as age, gender, education, ADNI cohort, and scanner manufacturer were included to adjust for different scan protocols and other potential scanner-related differences. All 2x2 (region x CSF group) mixed ANCOVAs were carried out in IBM SPSS statistics version 25 (SPSS) with type III sums of squares, and within-subject effects were interpreted without covariates ^32^.

#### Linear regression of disease status based on CSF marker

To better understand the relationship between disease status and functional MRI properties, CSF ratios (see section CSF biomarker) and functional MRI signals were considered in a linear regression model in SPSS. The functional MRI signal served as dependent variable, and CSF ratio as independent variable. The regression was run with the z-scored data. Subsequently, the dependent overlapping correlations of NbM vs EC with CSF ratio were compared using cocor^33,34^.

#### Robust regression

To minimize the influence of outliers, especially in the APSC, robust regression models were carried out in MATLAB® R2020b with fitlm using the bisquare weight function with the default tuning constant. The same covariates as for the mixed ANCOVA were included in the model. Finally, the predictive models (NbM◊EC and EC◊NbM) were tested for each CSF group (normal and abnormal) and each functional property (fALFF and ReHo). The data was z-scored before entering the analysis to ensure comparability of the APSC and baseline signal.

#### Moderation analyses of independent samples

Moderation analyses were carried out in SPSS using the PROCESS macro ^35^ for fALFF and ReHo investigating whether CSF group assignment moderates the spread (NbM◊EC vs. EC◊NbM) of functional degeneration. Here, CSF group was used as a dichotomous moderator variable. For the construction of products mean-centering was applied, and the heteroscedasticity consistent standard error HC3 (Davidson-MacKinnon) was applied.

#### Annual percentage signal change (APSC)

The following formula ^1,36^ was used to assess longitudinal APSC in fALFF and ReHo. It accounts for the days between both measurements and minimizes the influence of differences between both measurements within a subject.

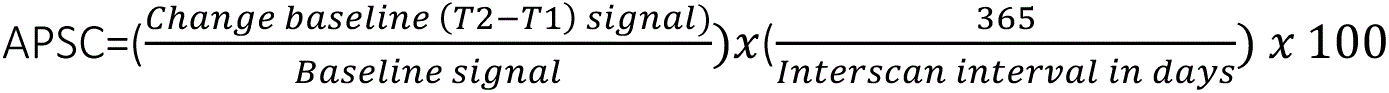

## Results

### CSF grouping strategy and neuropsychological assessments

Based on the CSF grouping strategy, we investigated how aCSF and nCSF groups performed in neuropsychological tests. For each test, one-way fixed effect ANOVAs were carried out with CSF group as factor and age, gender, and education as covariates. As expected, the nCSF group is less affected by cognitive impairment than the aCSF group (see Table 2).

### Lower fALFF values at baseline in aCSF vs. nCSF in NbM but not EC

Baseline fALFF values were compared in NbM and EC further subdivided into CSF groups using a 2x2 mixed ANCOVA. We found a main effect of CSF group (F(1,63)=7.943, p=0.006, partial η^2^=0.112, Fig. 1B), that was driven by lower fALFF values in participants with abnormal CSF, and a significant region x CSF group interaction (F(1,63)=4.623, p=0.035, partial η^2^=0.068, Fig. 1B), that was driven by a more pronounced fALFF reduction in the NbM. Post hoc analyses showed that a significant difference in fALFF between nCSF vs aCSF was only observed in NbM (t(69)=3.141, p=0.002) but not EC (t(69)=1.856, p=0.068). There was no main effect of region (F(1,69)=2.643, p=0.109, partial η^2^=0.037, Fig. 1B).

**Figure 1.**
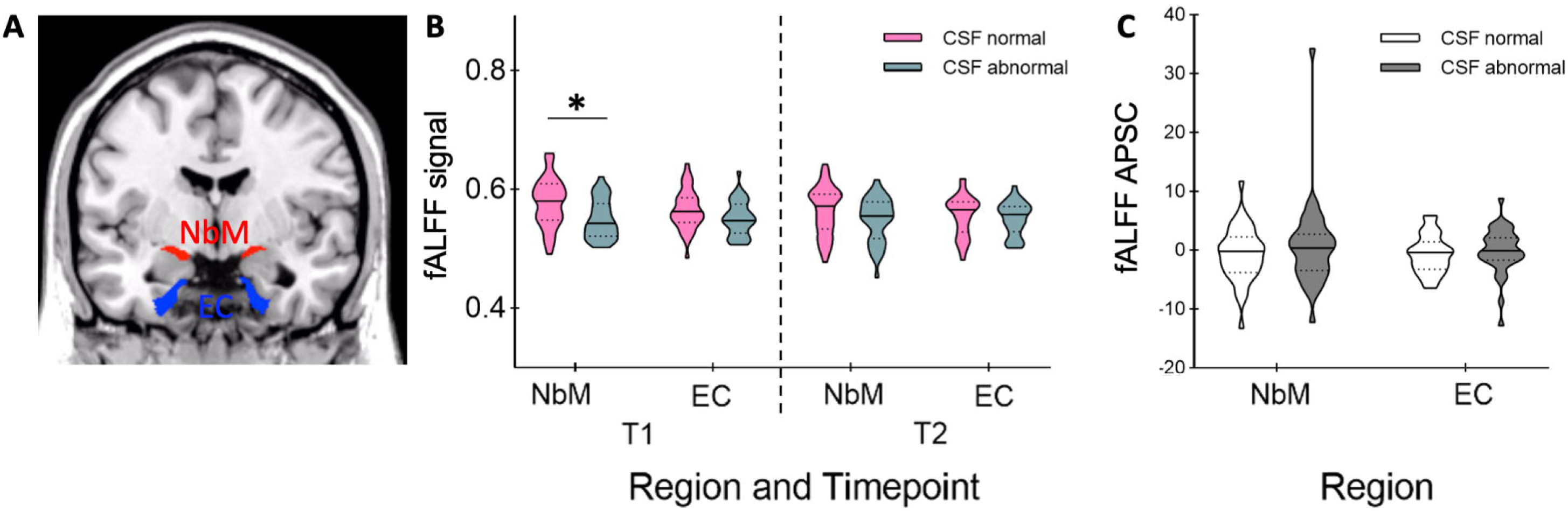
Region of interests, baseline signal and annual percentage signal change for fALFF measures. A) ROIs of NbM (red) and EC (blue) on a coronal slice of a T1-weighted standard brain template. Violin plots representing B) the baseline signals at time point 1 (T1) and the signals at the follow-up measurement (T2). C) shows the annual percentage signal change (APSC) in both regions. The horizontal lines represent medians and dotted lines interquartile ranges. *p<0.01.

### Annual percentage signal change in fALFF does not differentiate between CSF groups or regions

We used a 2x2 mixed ANCOVA to investigate whether the longitudinal indices of APSC in fALFF of the NbM and EC differentiated between CSF normal vs. abnormal groups. There was no significant main effect of CSF group (F(1,63)=2.077, p=0.154, partial η^2^=0.032, Fig. 1C), or region (F(1,69)=0.499, p=0.482, partial η^2^=0.007, Fig. 1C), and no significant group x region interaction (F(1,63)=0.367, p=0.547, partial η^2^=0.006, Fig. 1C) in APSC fALFF.

### NbM’s fALFF relates to CSF ratio

In a next step, we used linear regressions on fALFF values from NbM and EC, respectively, with CSF ratio as independent variable. It revealed a significant linear effect in the NbM (R²=0.120, F(1, 69)=9.437, p=0.003, Fig. 2A) but not EC (R²=0.031, F(1, 69)=2.206, p=0.142, Fig. 2B). A direct comparison of both correlations (NbM vs EC, one-tailed, which was justified by our a priori hypotheses) revealed a significant difference that was driven by a more negative correlation in NbM compared to EC (z=-1,94; p=0.0262; 95% CI: -0.3429 to 0.0015).

**Figure 2.**
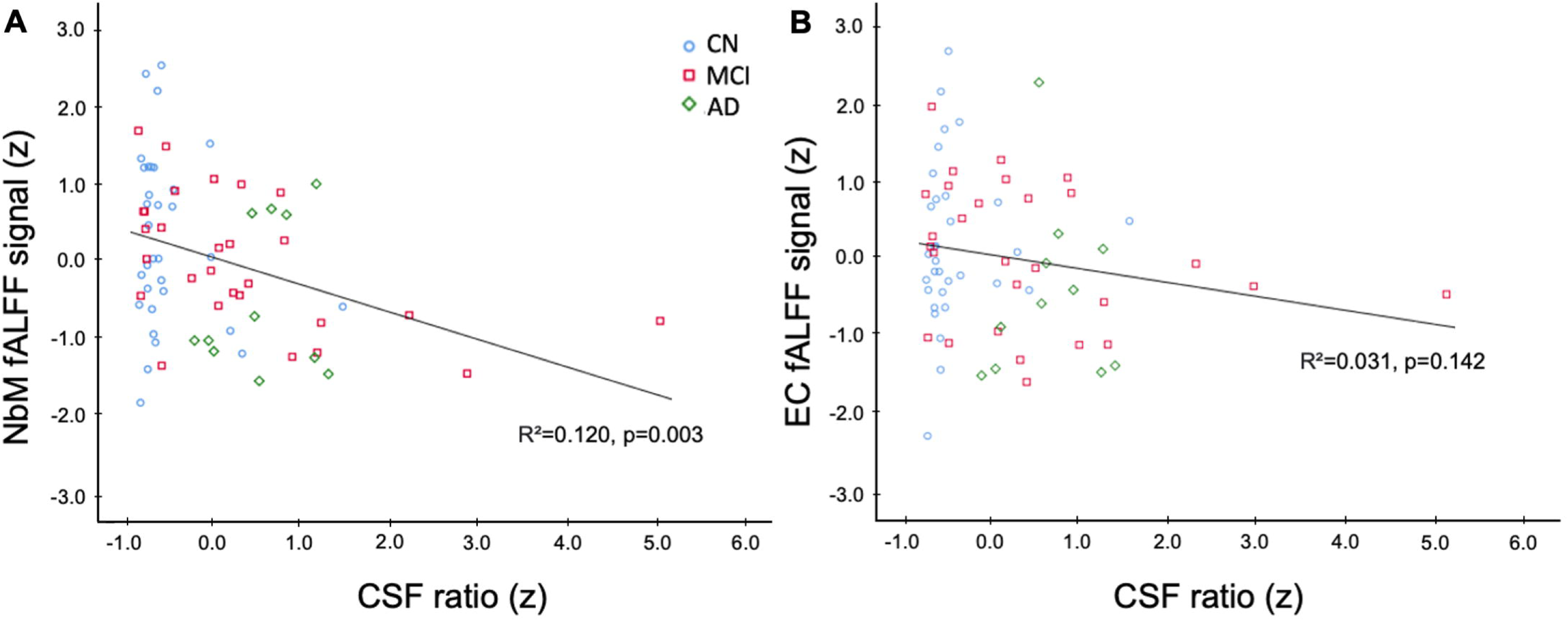
Linear regression for z-scored fALFF signal at baseline in A) NbM and B) EC against the z-scored CSF ratio. For the sake of visualization, the diagnosis groups are plotted in different colors and shapes (blue circle for CN, red square for MCI and green rhombus for AD). A significant linear regression was observed only in the NbM (A) but not EC (B), indicating a region specific decrease in functional activity and proteinopathology.

### Baseline signal in NbM predicts annual percentage signal change in fALFF of EC

To further address the temporal changes in AD progression, we examined whether the baseline signal in one region predicts the APSC in the other region. Here, in a first step, we used robust regression modeling for both competing models separately for nCSF vs. aCSF. They revealed no significant effect for NbM◊EC in aCSF (R²=0.263, F(7, 26)=1.33, p=0.277, Fig. 3A, Table S1), and no significant effect for NbM◊EC in nCSF (R²=0.296, F(7, 29)=1.74, p=0.138, Fig. 3B, Table S1). Similarly, there was no significant effect for EC◊NbM in aCSF (R²=0.137, F(7, 26)=0.587, p=0.76, Fig. 3D, Table S1), and no significant effect for EC◊NbM in nCSF (R²=0.175, F(7, 29)=0.88, p=0.534, Fig. 3E, Table S1).

**Figure 3.**
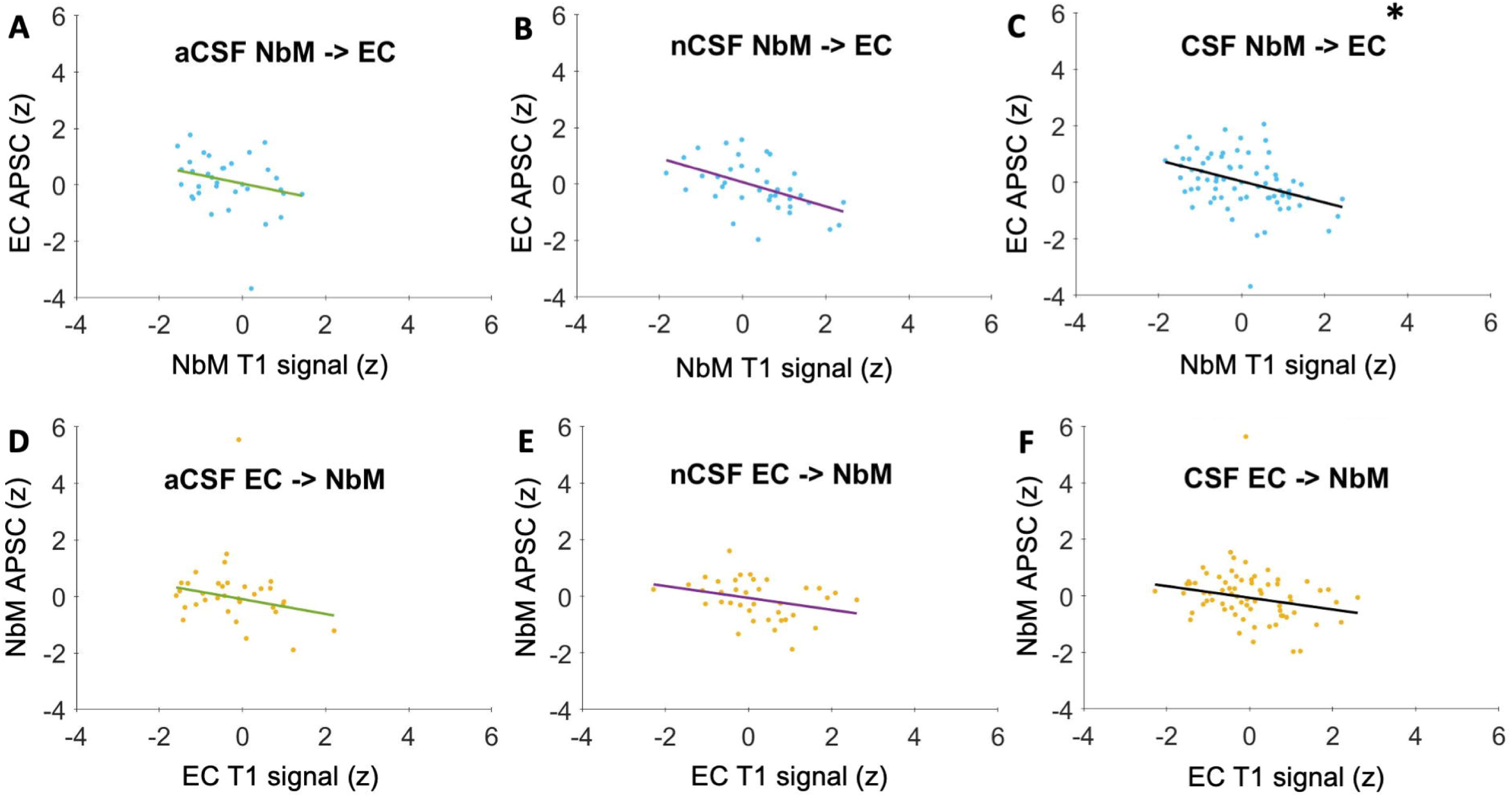
Adjusted for covariates response plot for the robust regression models in fALFF A) – C) predictive spread in NbM◊EC and D) – F) predictive spread in EC◊NbM. On the x-axis z-scores for the baseline signal in the respective region, on the y-axis the z-scored annual percentage signal change (APSC). There was a significant spread in C) NbM◊EC with CSF included in the model. There was no predictive spread in EC◊NbM. *p<0.05.

In a second step, we analyzed both groups together by including CSF group in the two competing regression models. Importantly, we observed a statistically significant effect for the model NbM◊EC (R²=0.235, F(8, 62)=2.39, p=0.026, Fig. 3C, Table S1), with NbM as a significant predictor of EC’s APSC (r=-0.3751, t(62)=-3.1445, p=0.003, confidence interval (CI): -0.6136 to 0.1366). The other regression model EC◊NbM did not show a significant effect (R²= 0.0884, F(8, 62)= 0.751, p= 0.646, Fig. 3F, Table S1). Replacing CSF as dichotomous predictor by the continuous CSF ratio did not change the results (i.e. significant effects for the model NbM◊EC, p=0.021, but not EC◊ NbM, p=0.466).

### CSF group does not moderate the relationship of NbM and EC in fALFF

Finally, we performed two moderation analyses. The first model included baseline fALFF NbM as independent variable, fALFF EC APSC as dependent variable and CSF group as moderator. The model was statistically significant (R²=0.2215, F(9,61)=3.4009, p=0.0019), with a significant direct effect of NbM◊EC (t(61)=-3.4420, p=0.001), but, no significant moderator effect (t(61)=0.4095, p=0.6836), which is in line with the robust regression analysis.

The second model included baseline fALFF EC as independent variable, fALFF NbM APSC as dependent variable and CSF group as moderator. The model was not statistically significant (R²=0.0965, F(9,61)=0.7857, p=0.6303), which, again, is in line with the robust regression analysis.

The results for ReHo and functional connectivity can be found in the supplementary material.

## Discussion

We investigated the functional properties of the human NbM and EC in relation to the disease progression of AD based on longitudinal rsfMRI data and CSF markers of Aβ and pTau. With a focus on fALFF, our data provide evidence that spontaneous local brain activity in the NbM, but not EC, is reduced with CSF ratio, and, importantly, it predicts the annual percentage signal change in the interconnected EC independently from proteinopathology. As such, our findings extend previous anatomical studies in humans and animals by providing novel physiological insights into the pathological staging model of AD suggesting the NbM as origin for subsequently affected brain regions possibly via a trans-synaptic mechanism.

Local spontaneous brain activity, as quantified by fALFF, was reduced in the NbM at baseline in the abnormal CSF group (Fig. 1B), and there was a linear reduction in fALFF activity with CSF-ratio (Fig. 2A). Importantly, both relationships were only observed in the NbM but not in the EC (Fig. 2B), which further underlines that the NbM is specifically vulnerable to AD progression. In fact, pTau and Aβ are two proteins that have been associated with AD ^37^ and the NbM is particularly vulnerable to the early accumulation of pTau ^38–40^ and Aβ deposition ^41^. This may be due to the fact that cholinergic basal forebrain neurons have rather large axons and arbors reaching into the entire central nervous system with high metabolic demands for maintenance, reparation, and transportation ^42^. At the same time, simply due to their sizes, they are more vulnerable to toxins ^43^, which may further promote disease progression.

The pathological staging model suggests a structural degeneration originating in the NbM followed by the EC, which adds a crucial upstream link to Alzheimer’s degeneration ^1^. Our functional data support such a view by showing that the NbM’s baseline fALFF signal predicted the APSC in the EC (Figure 3C) but not the reverse (Figure 3F). Interestingly, this effect was independent of CSF status, which was further supported by the absence of a moderating effect of CSF. While this is compatible with a specific spread from NbM to EC, it also indicates that the putative functional consequences, namely changes in neural activation, are unrelated to pTau and Aβ. This apparently differs from anatomical changes from NbM to EC that were more pronounced in subjects with abnormal CSF ^1^. From a physiological point of view, a trans-synaptic spread of proteins between anatomically interconnected brain regions is possible and has been shown in several animal studies. For instance, aggregates of tau can propagate from the EC to other limbic regions, including the dentate gyrus and hippocampal CA fields, followed by neocortical brain regions including the parietal cortex ^3–5,44^. *In vitro*, this can be enhanced by neural activity ^6^, which might help to explain why CSF status did not moderate the relationship between NbM activity and longitudinal changes in EC activity in our study. While this needs to be further investigated using larger and independent samples, our study is the first to show *in vivo* in humans that a neural signal in NbM can serve as a predictive marker for functional changes in the anatomically interconnected EC across healthy controls, MCIs and AD patients.

Although fALFF is a prominent marker of spontaneous local brain activity ^11,12^, only a limited number of studies used fALFF to investigate AD. Importantly, previous work did not specifically focus on the NbM and EC but other, typically larger brain regions. It showed, for instance, decreased fALFF signals in the bilateral middle frontal and left precuneus in participants with positive Aβ ^14^. In preclinical AD, increases and decreases in fALFF were reported in the right inferior frontal gyrus ^14,45^, and in prodromal AD lower fALFF signals could be shown in the bilateral precuneus, right middle frontal gyrus, right precentral gyrus, and postcentral gyrus. Finally, in AD fALFF was increased in the right fusiform gyrus, medial temporal lobe and inferior temporal gyrus, but decreased in the bilateral precuneus, left posterior cingulate cortex, left cuneus and superior occipital gyrus ^45^. These partly divergent effects of fALFF associated with AD might be explained by compensatory effects to maintain an adequate level of cognitive performance ^45^, and could be a functional hallmark of neural aging ^46^ that needs further attention. Furthermore, since no significant effects in ReHo and functional connectivity between were detected (see supplementary material), fALFF seems to be a particularly sensitive marker. Together, fALFF is highly sensitive to changes in neural activity associated with AD even in rather small brain regions and therefore offers a useful marker in future studies.

Our analyses specifically focused on the functional properties of the human NbM and EC but no other interconnected brain regions that, according to the pathological staging model, follow the EC. These may include the parahippocampal cortex and hippocampal structures, as well as the parietal cortex ^1,3–5,44^. Along these lines, we included functional signals averaged from both hemispheres, which simplified our analyses, but it neglected possible lateralization effects ^47,48^. Second, ADNI is a large multicenter study offering a rich and unique dataset. However, our rsfMRI data come from different MR scanners, possibly leading to a bias in image quality and extracted signal. Therefore, we only included high-quality data that were based on comparable protocols and within-subject measurements from the same scanner. We also employed appropriate covariates in our statistical models, and differences in scanning parameters (e.g. slice order or number of volumes) were accounted for by during preprocessing ^49,50^. Further, our main findings are based on analyses including a measure of APSC, which is robust against within-subject variability, e.g., because of the MR scanner.

Functional activity in the human basal forebrain decreased with proteinopathology and predicted the functional decrease within the interconnected EC independent from CSF status. As such, our findings extend the pathological staging model of AD by giving novel insights into the functional properties of the underlying brain regions. From a more general perspective, fALFF appears to be a suitable marker to further investigate functional brain changes associated with the progression of AD.

## Acknowledgments

Data collection and sharing for this project was funded by the Alzheimer’s Disease Neuroimaging Initiative (ADNI) (National Institutes of Health Grant U01 AG024904) and DOD ADNI (Department of Defense award number W81XWH-12-2-0012). ADNI is funded by the National Institute on Aging, the National Institute of Biomedical Imaging and Bioengineering, and through generous contributions from the following: AbbVie, Alzheimer’s Association; Alzheimer’s Drug Discovery Foundation; Araclon Biotech; BioClinica, Inc.; Biogen; Bristol-Myers Squibb Company; CereSpir, Inc.; Cogstate; Eisai Inc.; Elan Pharmaceuticals, Inc.; Eli Lilly and Company; EuroImmun; F. Hoffmann-La Roche Ltd and its affiliated company Genentech, Inc.; Fujirebio; GE Healthcare; IXICO Ltd.; Janssen Alzheimer Immunotherapy Research & Development, LLC.; Johnson & Johnson Pharmaceutical Research & Development LLC.; Lumosity; Lundbeck; Merck & Co., Inc.; Meso Scale Diagnostics, LLC.; NeuroRx Research; Neurotrack Technologies; Novartis Pharmaceuticals Corporation; Pfizer Inc.; Piramal Imaging; Servier; Takeda Pharmaceutical Company; and Transition Therapeutics. The Canadian Institutes of Health Research is providing funds to support ADNI clinical sites in Canada. Private sector contributions are facilitated by the Foundation for the National Institutes of Health (www.fnih.org). The grantee organization is the Northern California Institute for Research and Education, and the study is coordinated by the Alzheimer’s Therapeutic Research Institute at the University of Southern California. ADNI data are disseminated by the Laboratory for Neuro Imaging at the University of Southern California.

---

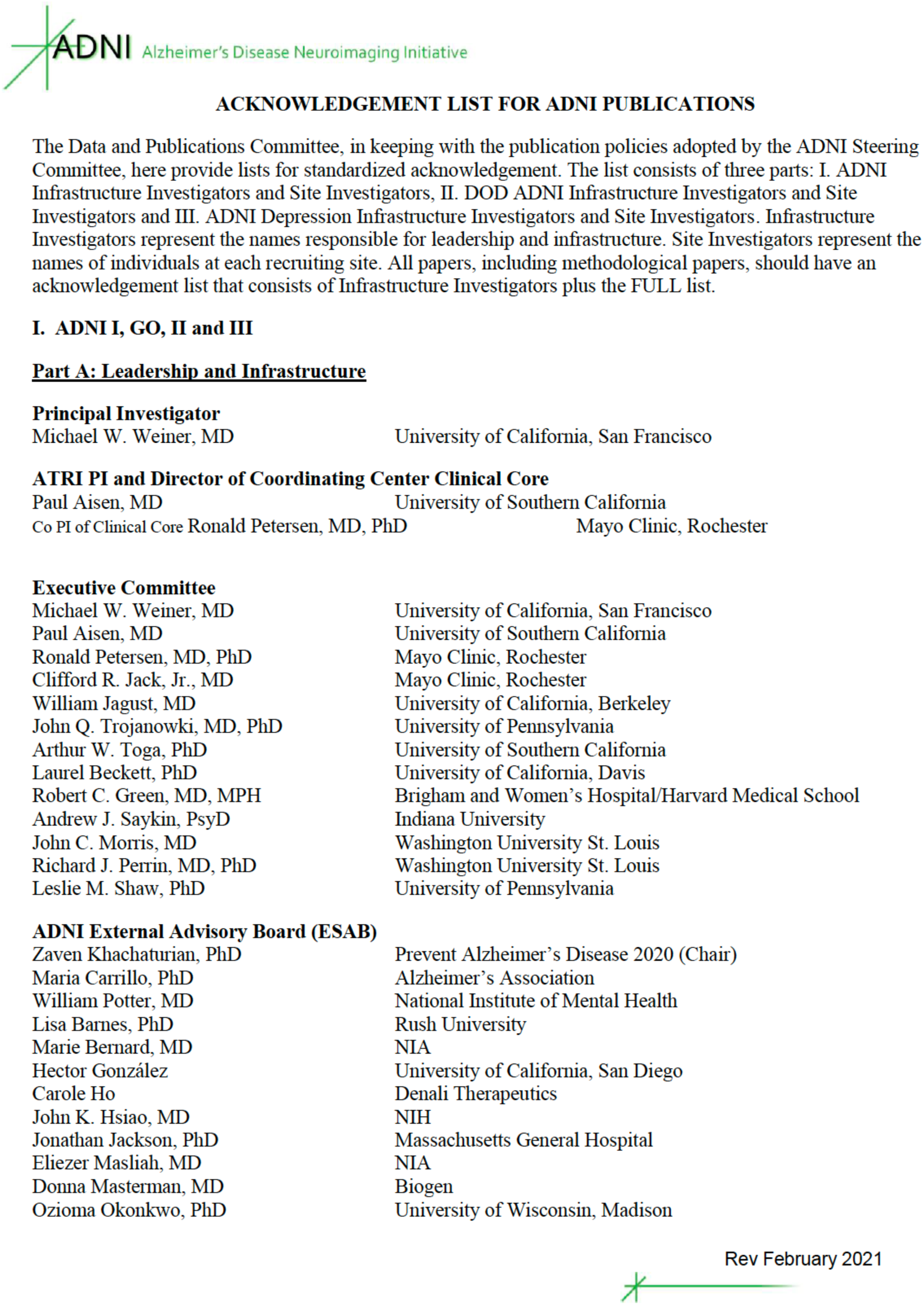

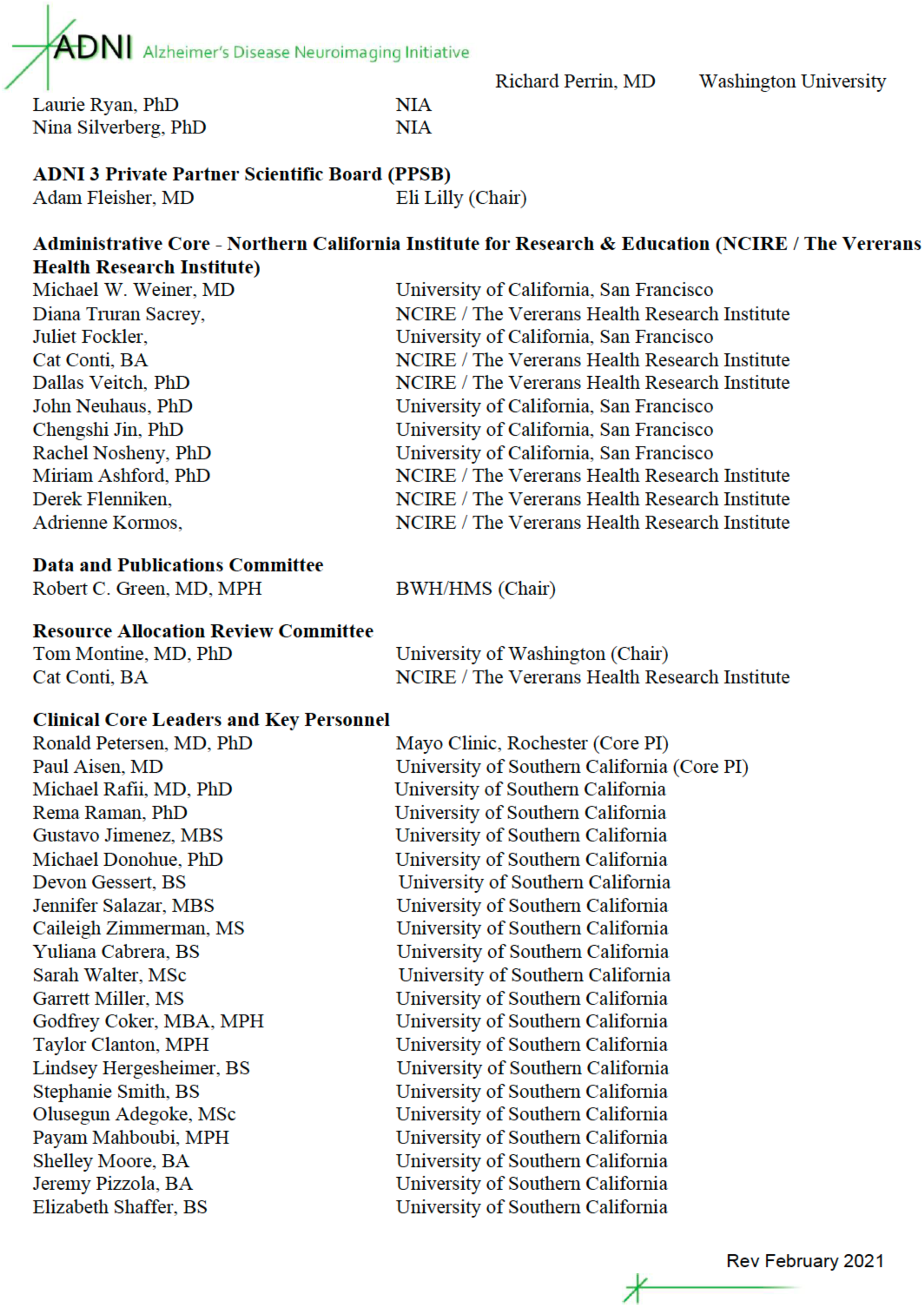

|  |  |
| --- | --- |
| Michael W. Weiner, MD | University of California, San Francisco |
| Diana Truran Sacrey, | NCIRE / The Veterans Health Research Institute |
| Juliet Fockler, | University of California, San Francisco |
| Cat Conti, BA | NCIRE / The Veterans Health Research Institute |
| Dallas Veitch, PhD | NCIRE / The Veterans Health Research Institute |
| John Neuhaus, PhD | University of California, San Francisco |
| Chengshi Jin, PhD | University of California, San Francisco |
| Rachel Nosheny, PhD | University of California, San Francisco |
| Miriam Ashford, PhD | NCIRE / The Veterans Health Research Institute |
| Derek Flenniken, | NCIRE / The Veterans Health Research Institute |
| Adrienne Kormos, | NCIRE / The Veterans Health Research Institute |
| **Resource Allocation Review Committee** |  |
| Tom Montine, MD, PhD | University of Washington (Chair) |
| Cat Conti, BA | NCIRE / The Veterans Health Research Institute |
| **Clinical Core Leaders and Key Personnel** |  |
| Ronald Petersen, MD, PhD | Mayo Clinic, Rochester (Core PI) |
| Paul Aisen, MD | University of Southern California (Core PI) |
| Michael Rafii, MD, PhD | University of Southern California |
| Rema Raman, PhD | University of Southern California |
| Gustavo Jimenez, MBS | University of Southern California |
| Michael Donohue, PhD | University of Southern California |
| Devon Gessert, BS | University of Southern California |
| Jennifer Salazar, MBS | University of Southern California |
| Caileigh Zimmerman, MS | University of Southern California |
| Yuliana Cabrera, BS | University of Southern California |
| Sarah Walter, MSc | University of Southern California |
| Garrett Miller, MS | University of Southern California |
| Godfrey Coker, MBA, MPH | University of Southern California |
| Taylor Clanton, MPH | University of Southern California |
| Lindsey Hergesheimer, BS | University of Southern California |
| Stephanie Smith, BS | University of Southern California |
| Olusegun Adegoke, MSc | University of Southern California |
| Payam Mahboubi, MPH | University of Southern California |
| Shelley Moore, BA | University of Southern California |
| Jeremy Pizzola, BA | University of Southern California |
| Elizabeth Shaffer, BS | University of Southern California |

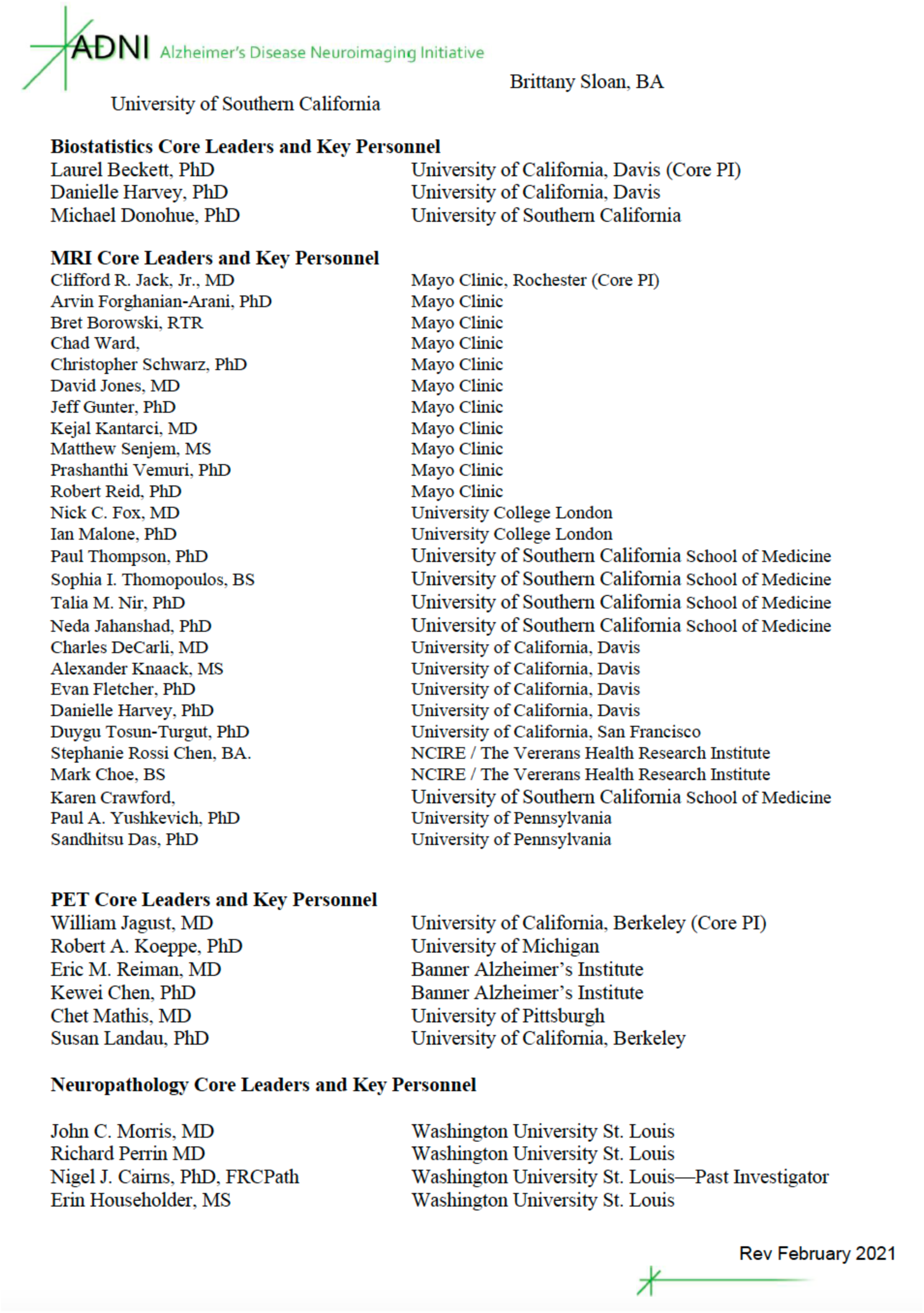

Biostatistics Core Leaders and Key Personnel
|  |  |
| --- | --- |
| Laurel Beckett, PhD | University of California, Davis (Core PI) |
| Danielle Harvey, PhD | University of California, Davis |
| Michael Donohue, PhD | University of Southern California |
| MRI Core Leaders and Key Personnel |  |
| Clifford R. Jack, Jr., MD | Mayo Clinic, Rochester (Core PI) |
| Arvin Forghanian-Arani, PhD | Mayo Clinic |
| Bret Borowski, RTR | Mayo Clinic |
| Chad Ward, | Mayo Clinic |
| Christopher Schwarz, PhD | Mayo Clinic |
| David Jones, MD | Mayo Clinic |
| Jeff Gunter, PhD | Mayo Clinic |
| Kejal Kantarci, MD | Mayo Clinic |
| Matthew Senjem, MS | Mayo Clinic |
| Prashanthi Vemuri, PhD | Mayo Clinic |
| Robert Reid, PhD | Mayo Clinic |
| Nick C. Fox, MD | University College London |
| Ian Malone, PhD | University College London |
| Paul Thompson, PhD | University of Southern California School of Medicine |
| Sophia I. Thomopoulos, BS | University of Southern California School of Medicine |
| Talia M. Nir, PhD | University of Southern California School of Medicine |
| Neda Jahanshad, PhD | University of Southern California School of Medicine |
| Charles DeCarli, MD | University of California, Davis |
| Alexander Knaack, MS | University of California, Davis |
| Evan Fletcher, PhD | University of California, Davis |
| Danielle Harvey, PhD | University of California, Davis |
| Duygu Tosun-Turgut, PhD | University of California, San Francisco |
| Stephanie Rossi Chen, BA. | NCIRE / The Veterans Health Research Institute |
| Mark Choe, BS | NCIRE / The Veterans Health Research Institute |
| Karen Crawford, | University of Southern California School of Medicine |
| Paul A. Yushkevich, PhD | University of Pennsylvania |
| Sandhitsu Das, PhD | University of Pennsylvania |
| PET Core Leaders and Key Personnel |  |
| William Jagust, MD | University of California, Berkeley (Core PI) |
| Robert A. Koeppe, PhD | University of Michigan |
| Eric M. Reiman, MD | Banner Alzheimer's Institute |
| Kewei Chen, PhD | Banner Alzheimer's Institute |
| Chet Mathis, MD | University of Pittsburgh |
| Susan Landau, PhD | University of California, Berkeley |
| Neuropathology Core Leaders and Key Personnel |  |
| John C. Morris, MD | Washington University St. Louis |
| Richard Perrin MD | Washington University St. Louis |
| Nigel J. Cairns, PhD, FRCPath | Washington University St. Louis—Past Investigator |
| Erin Householder, MS | Washington University St. Louis |

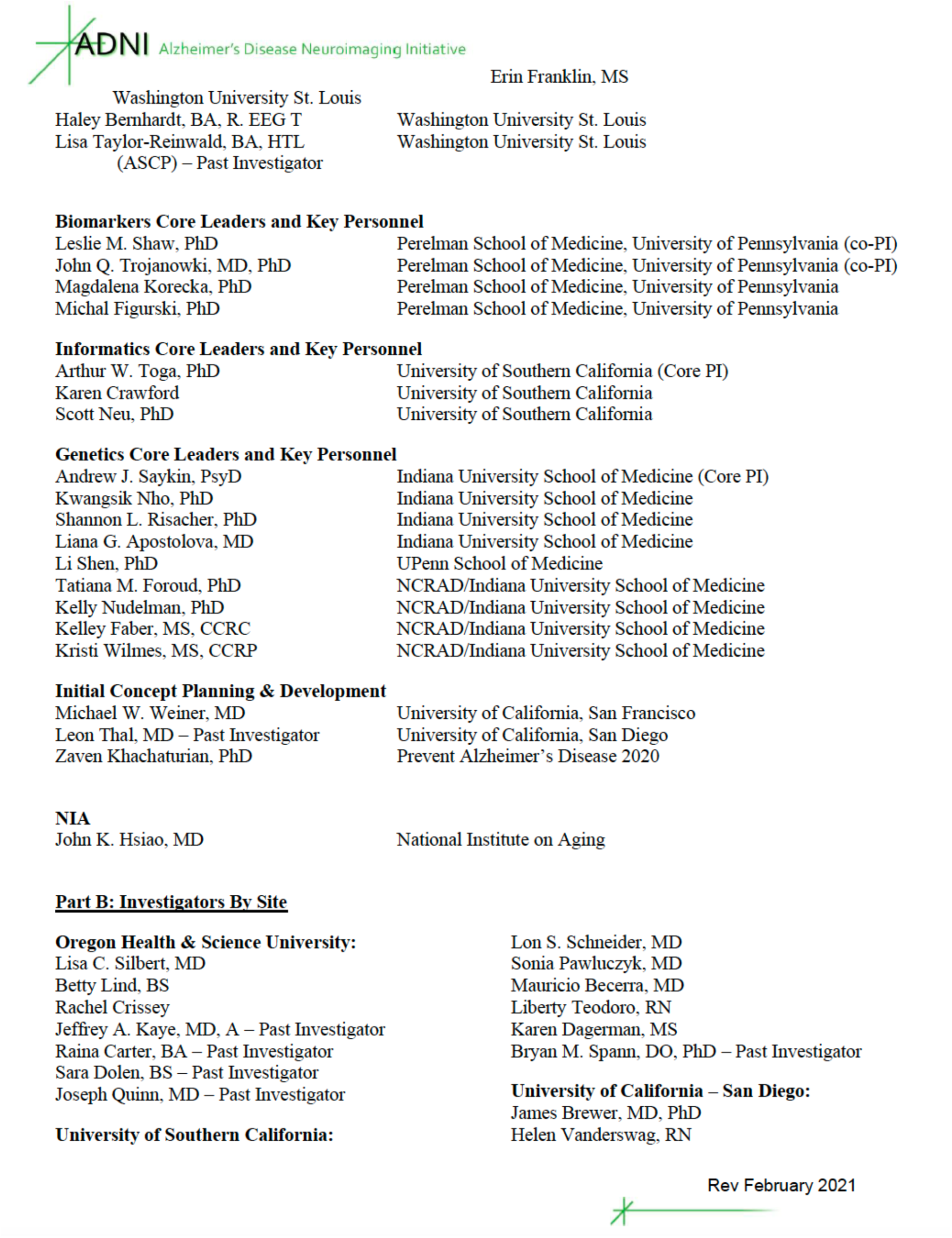

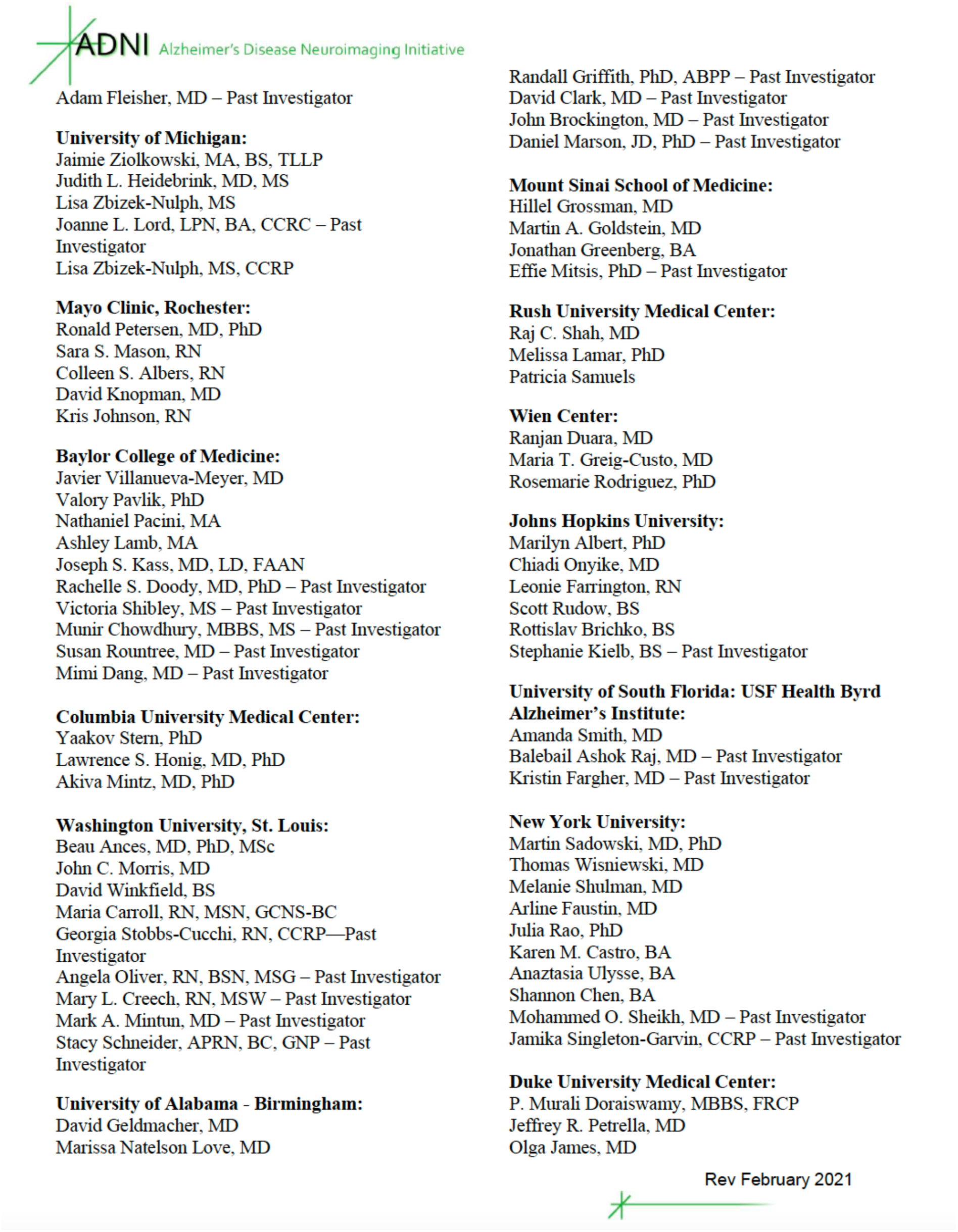

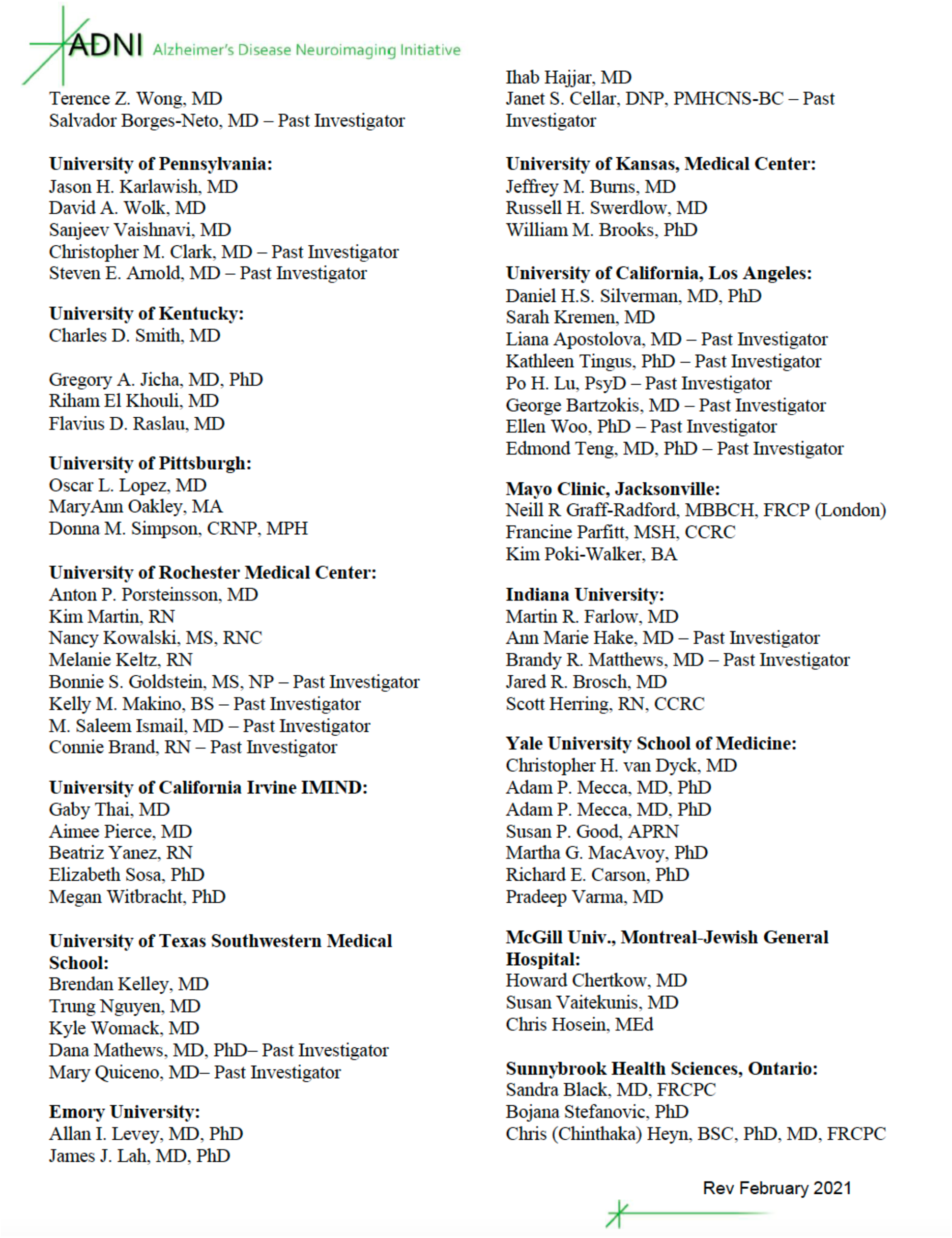

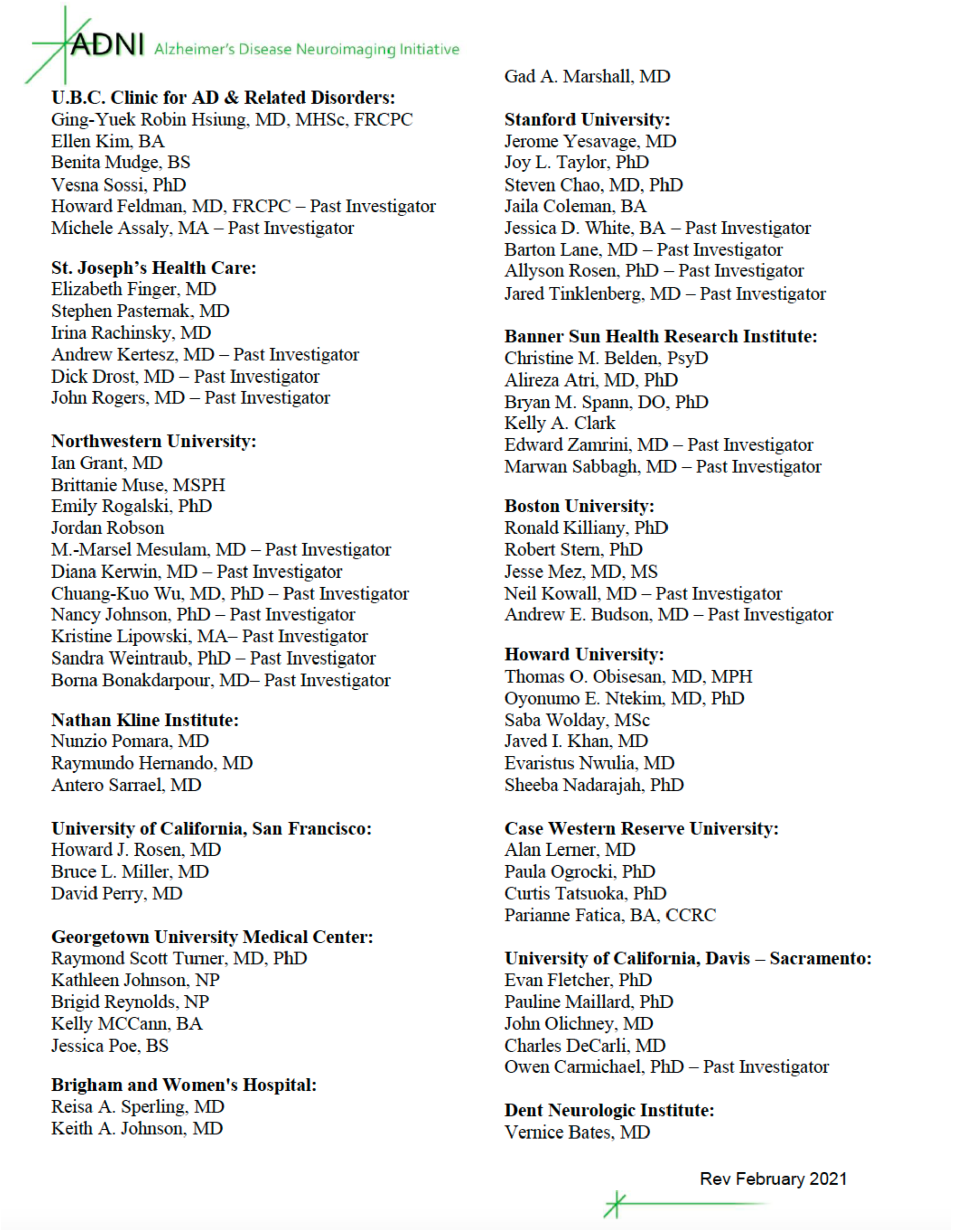

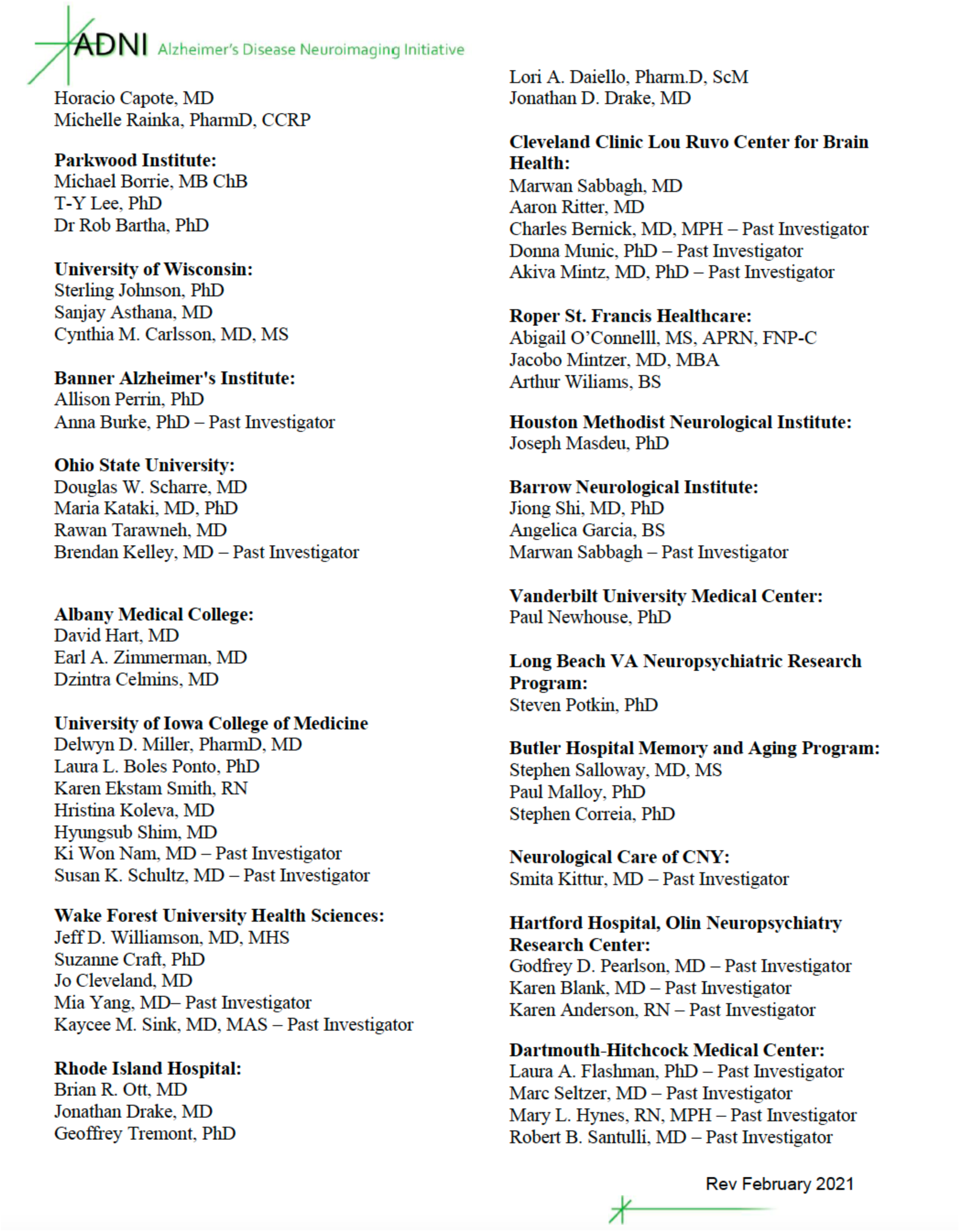

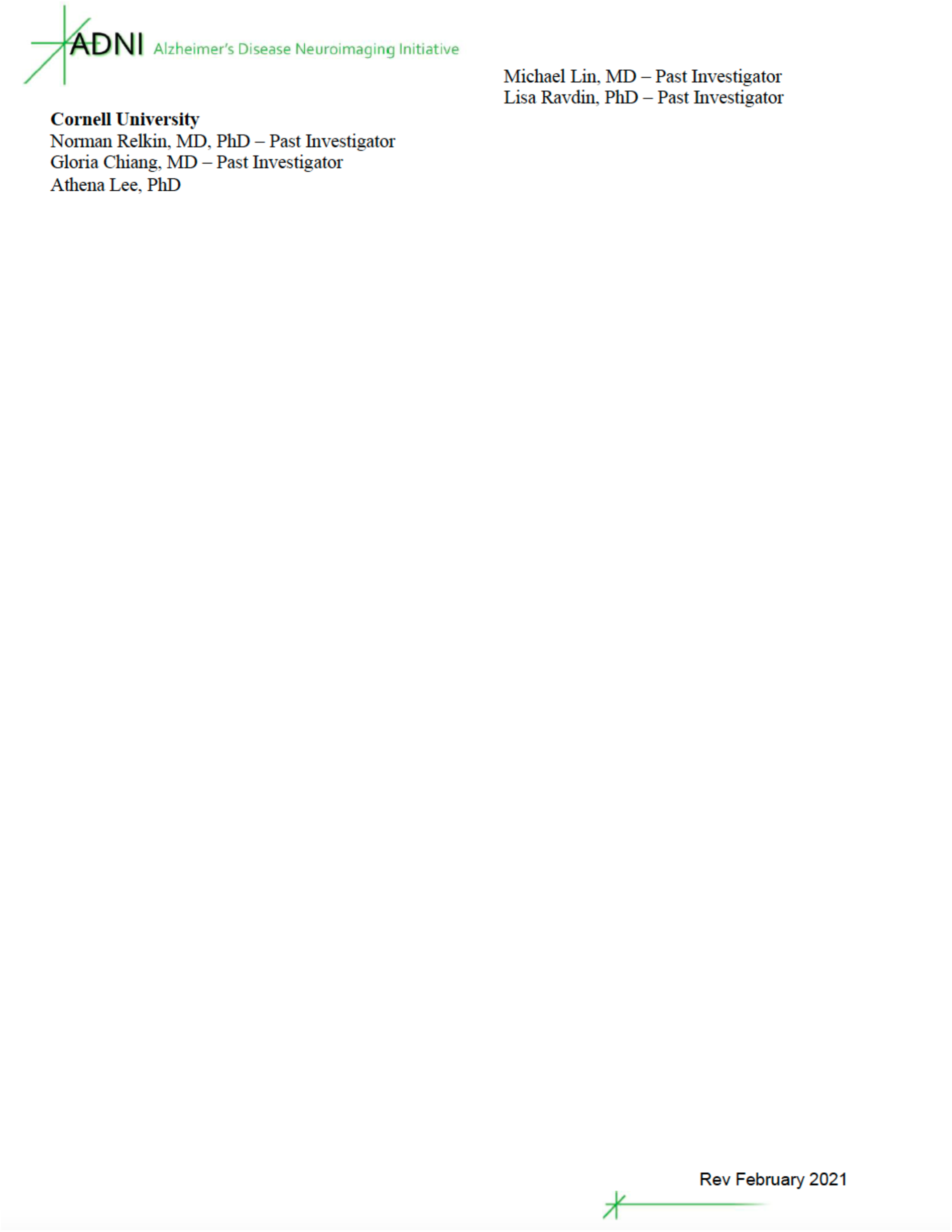

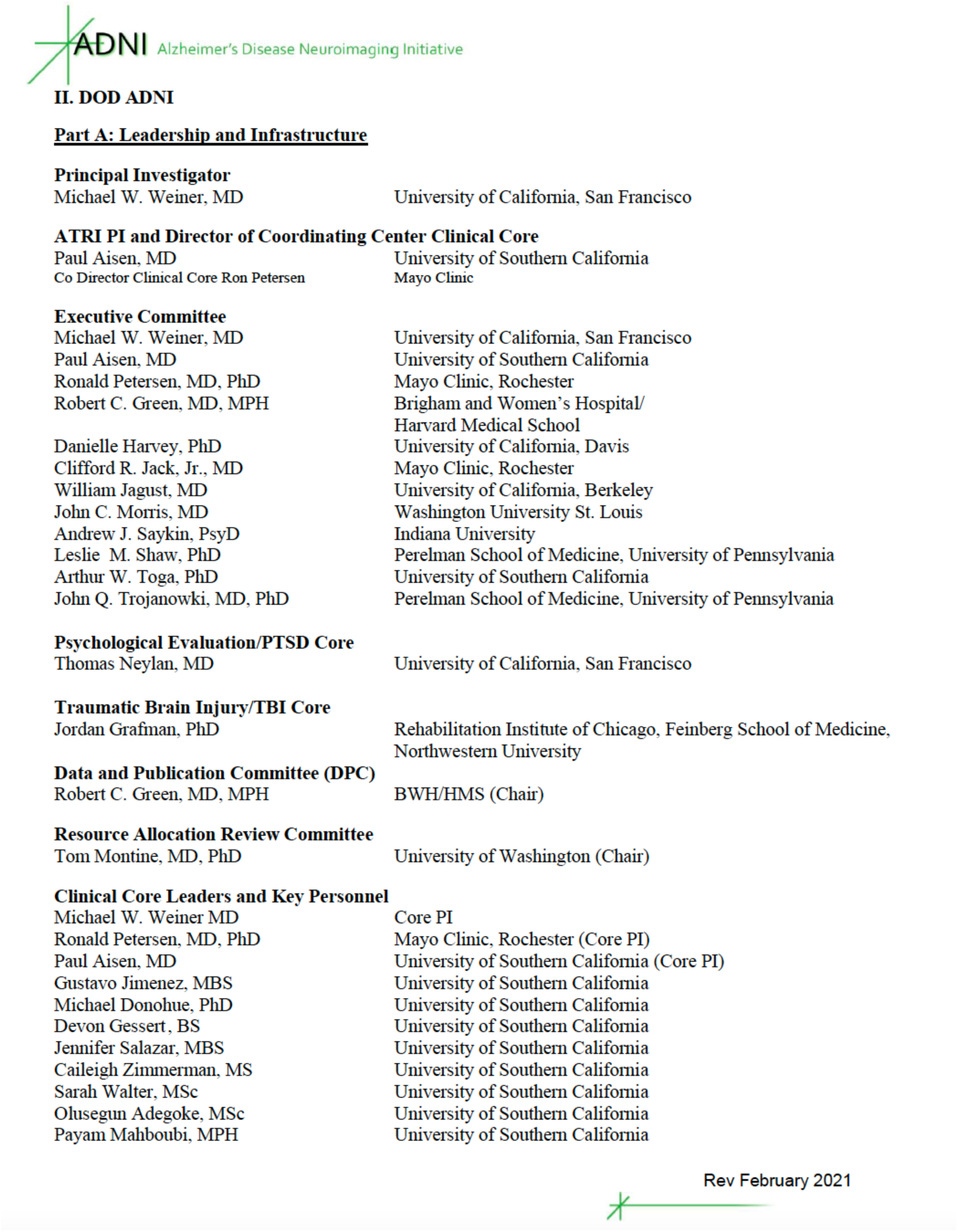

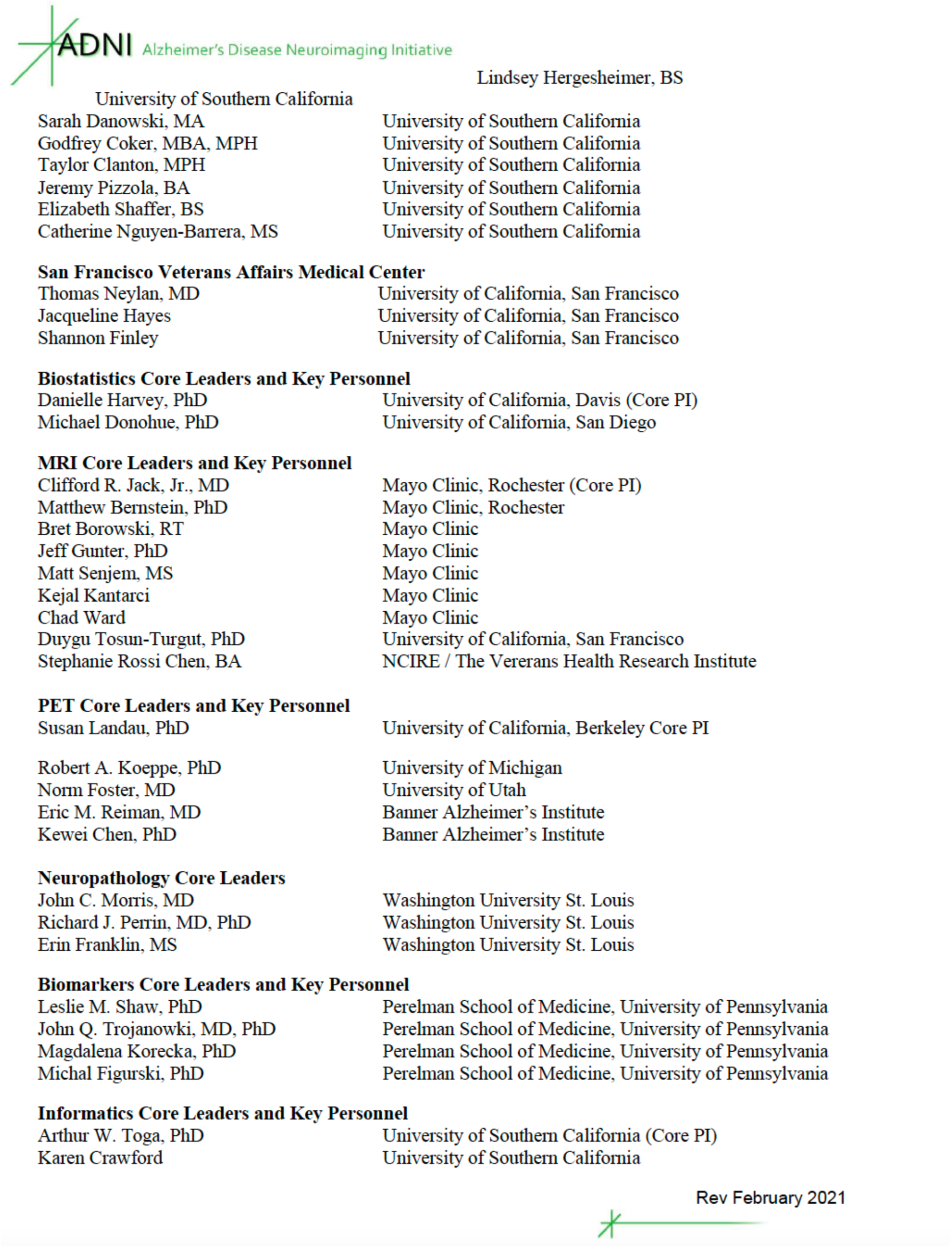

|  |  |
| --- | --- |
| Sarah Danowski, MA | University of Southern California |
| Godfrey Coker, MBA, MPH | University of Southern California |
| Taylor Clanton, MPH | University of Southern California |
| Jeremy Pizzola, BA | University of Southern California |
| Elizabeth Shaffer, BS | University of Southern California |
| Catherine Nguyen-Barrera, MS | University of Southern California |
| **San Francisco Veterans Affairs Medical Center** |  |
| Thomas Neylan, MD | University of California, San Francisco |
| Jacqueline Hayes | University of California, San Francisco |
| Shannon Finley | University of California, San Francisco |
| **Biostatistics Core Leaders and Key Personnel** |  |
| Danielle Harvey, PhD | University of California, Davis (Core PI) |
| Michael Donohue, PhD | University of California, San Diego |
| **MRI Core Leaders and Key Personnel** |  |
| Clifford R. Jack, Jr., MD | Mayo Clinic, Rochester (Core PI) |
| Matthew Bernstein, PhD | Mayo Clinic, Rochester |
| Bret Borowski, RT | Mayo Clinic |
| Jeff Gunter, PhD | Mayo Clinic |
| Matt Senjem, MS | Mayo Clinic |
| Kejal Kantarci | Mayo Clinic |
| Chad Ward | Mayo Clinic |
| Duygu Tosun-Turgut, PhD | University of California, San Francisco |
| Stephanie Rossi Chen, BA | NCIRE / The Veterans Health Research Institute |
| **PET Core Leaders and Key Personnel** |  |
| Susan Landau, PhD | University of California, Berkeley Core PI |
| Robert A. Koeppe, PhD | University of Michigan |
| Norm Foster, MD | University of Utah |
| Eric M. Reiman, MD | Banner Alzheimer's Institute |
| Kewei Chen, PhD | Banner Alzheimer's Institute |
| **Neuropathology Core Leaders** |  |
| John C. Morris, MD | Washington University St. Louis |
| Richard J. Perrin, MD, PhD | Washington University St. Louis |
| Erin Franklin, MS | Washington University St. Louis |
| **Biomarkers Core Leaders and Key Personnel** |  |
| Leslie M. Shaw, PhD | Perelman School of Medicine, University of Pennsylvania |
| John Q. Trojanowki, MD, PhD | Perelman School of Medicine, University of Pennsylvania |
| Magdalena Korecka, PhD | Perelman School of Medicine, University of Pennsylvania |
| Michal Figurski, PhD | Perelman School of Medicine, University of Pennsylvania |
| **Informatics Core Leaders and Key Personnel** |  |
| Arthur W. Toga, PhD | University of Southern California (Core PI) |
| Karen Crawford | University of Southern California |

Rev February 2021

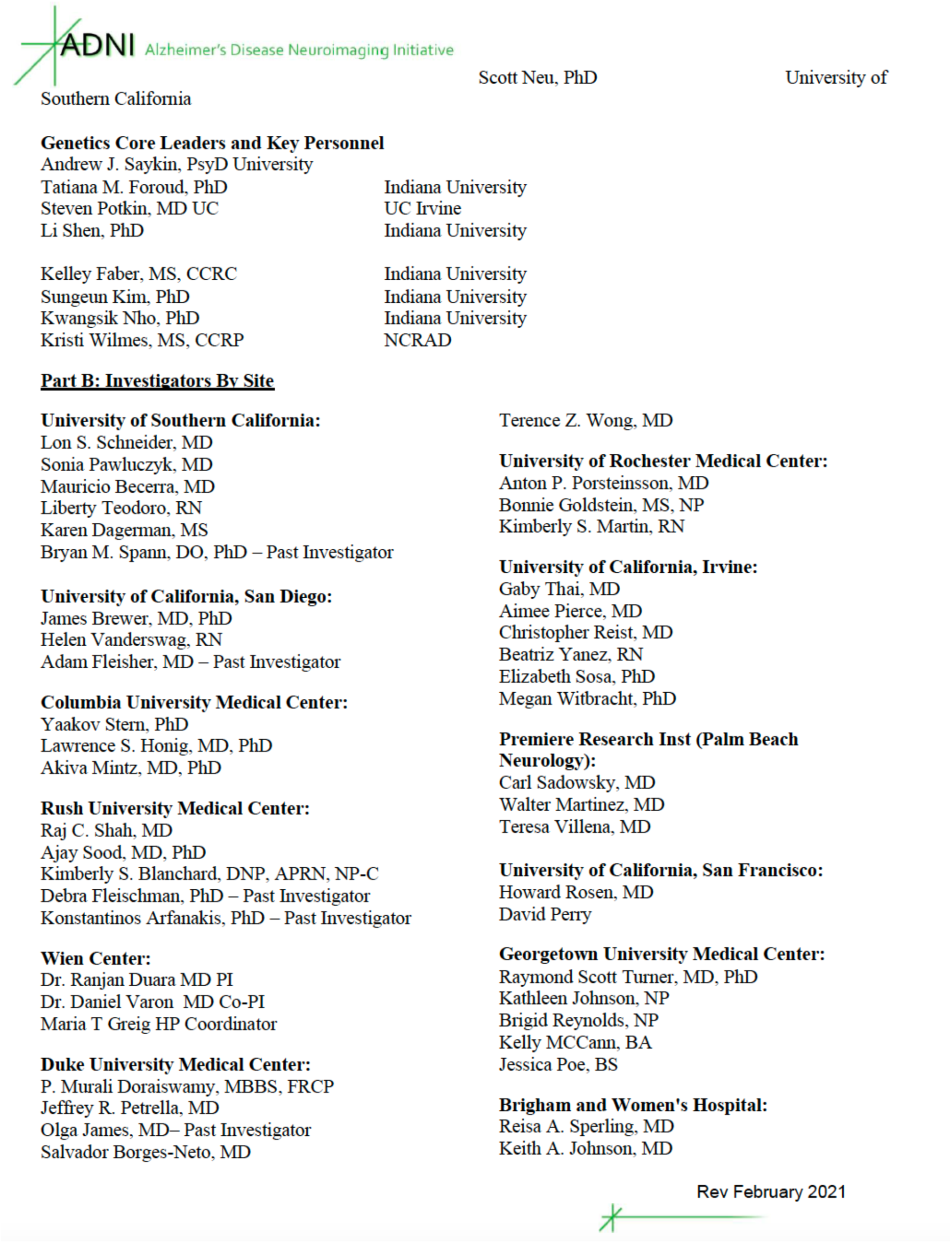

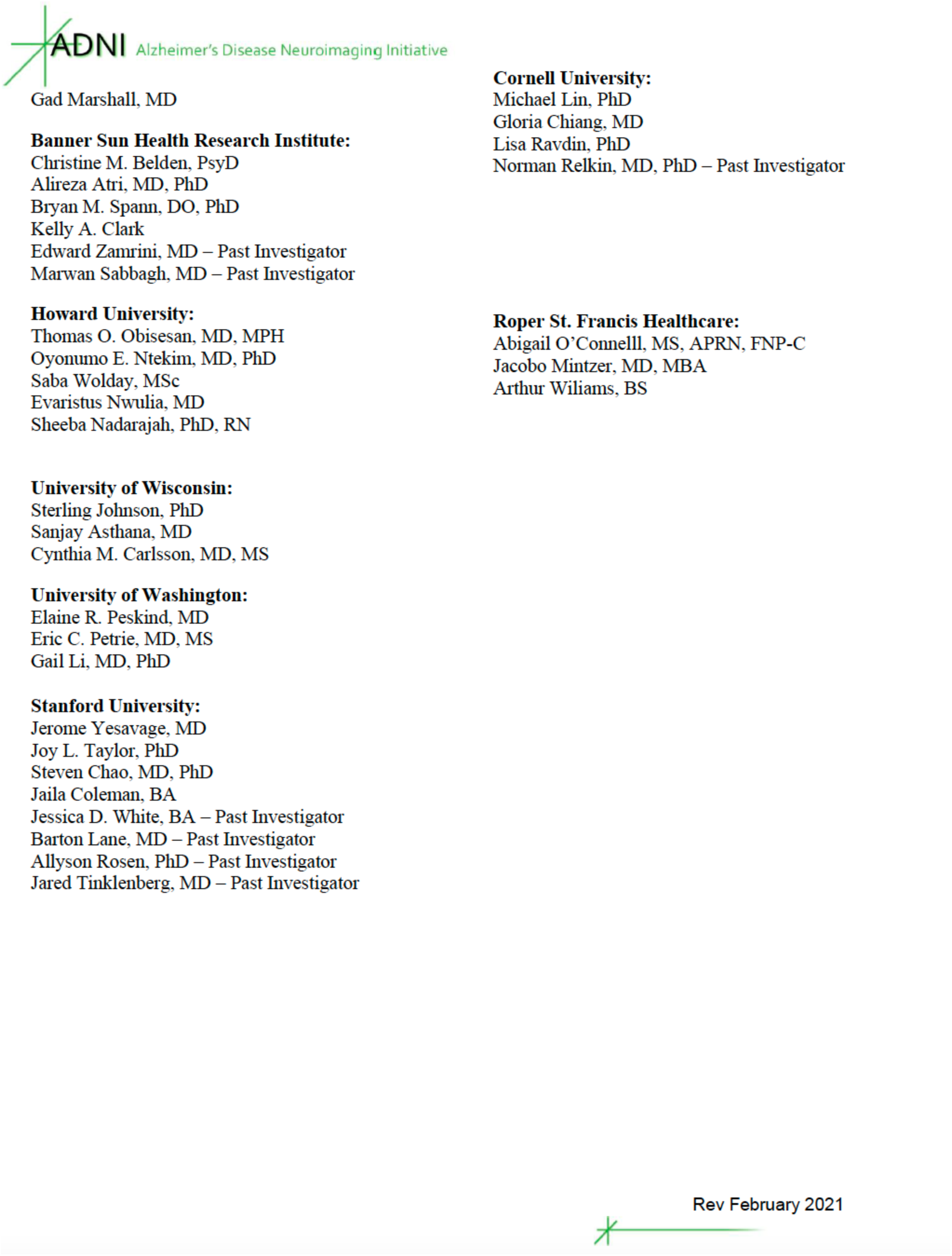

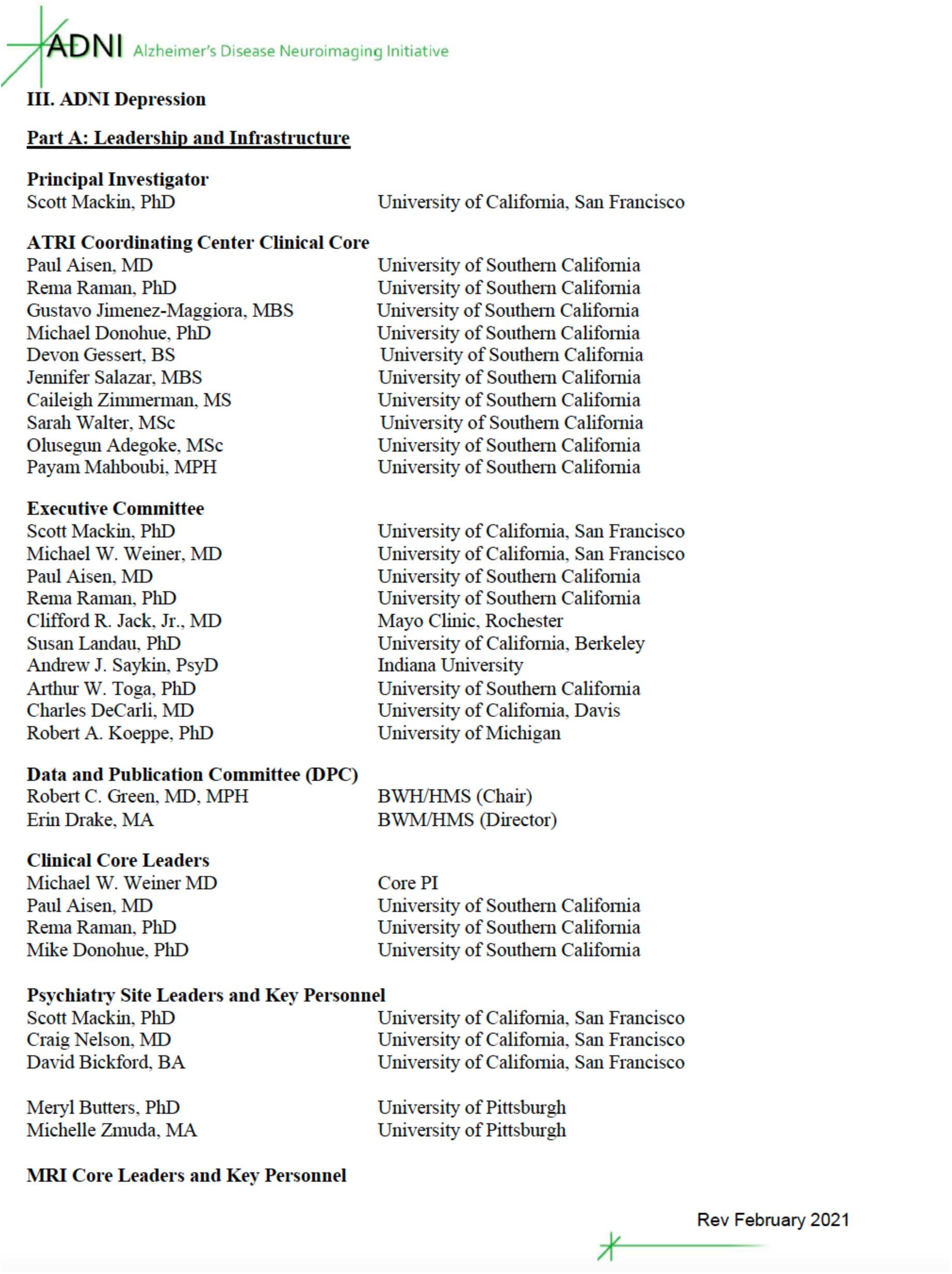

**ATRI Coordinating Center Clinical Core**
|  |  |
| --- | --- |
| Paul Aisen, MD | University of Southern California |
| Rema Raman, PhD | University of Southern California |
| Gustavo Jimenez-Maggiora, MBS | University of Southern California |
| Michael Donohue, PhD | University of Southern California |
| Devon Gessert, BS | University of Southern California |
| Jennifer Salazar, MBS | University of Southern California |
| Caileigh Zimmerman, MS | University of Southern California |
| Sarah Walter, MSc | University of Southern California |
| Olusegun Adegoke, MSc | University of Southern California |
| Payam Mahboubi, MPH | University of Southern California |
| **Executive Committee** |  |
| Scott Mackin, PhD | University of California, San Francisco |
| Michael W. Weiner, MD | University of California, San Francisco |
| Paul Aisen, MD | University of Southern California |
| Rema Raman, PhD | University of Southern California |
| Clifford R. Jack, Jr., MD | Mayo Clinic, Rochester |
| Susan Landau, PhD | University of California, Berkeley |
| Andrew J. Saykin, PsyD | Indiana University |
| Arthur W. Toga, PhD | University of Southern California |
| Charles DeCarli, MD | University of California, Davis |
| Robert A. Koeppe, PhD | University of Michigan |
| **Data and Publication Committee (DPC)** |  |
| Robert C. Green, MD, MPH | BWH/HMS (Chair) |
| Erin Drake, MA | BWH/HMS (Director) |
| **Clinical Core Leaders** |  |
| Michael W. Weiner MD | Core PI |
| Paul Aisen, MD | University of Southern California |
| Rema Raman, PhD | University of Southern California |
| Mike Donohue, PhD | University of Southern California |
| **Psychiatry Site Leaders and Key Personnel** |  |
| Scott Mackin, PhD | University of California, San Francisco |
| Craig Nelson, MD | University of California, San Francisco |
| David Bickford, BA | University of California, San Francisco |
| Meryl Butters, PhD | University of Pittsburgh |
| Michelle Zmuda, MA | University of Pittsburgh |

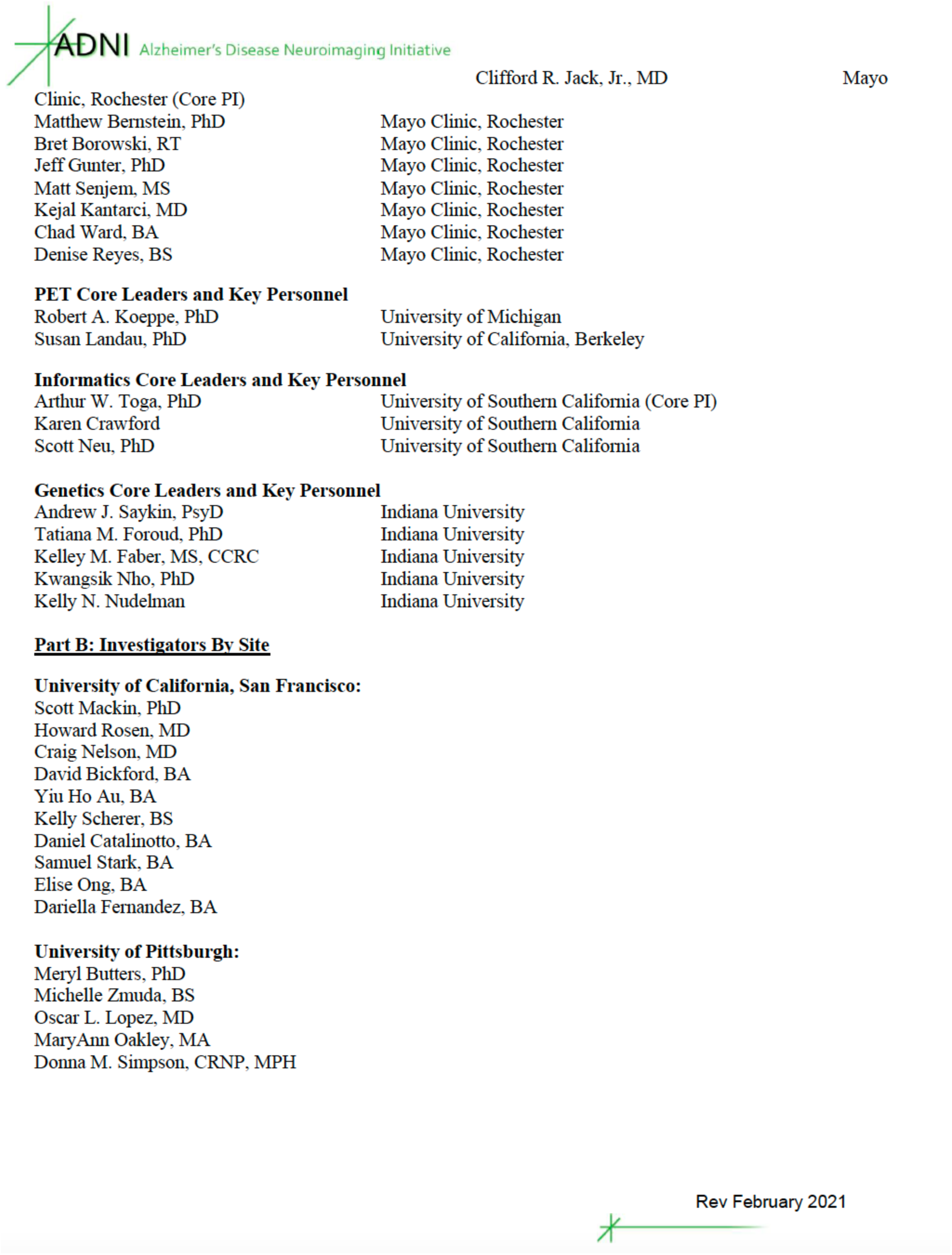

## Authors contributions

MM and NB conceived the study. MM, MG and MY analyzed the data. MM and NB wrote the manuscript. MG and MY gave further constructive feedback on the manuscript. All authors approved the manuscript.

## Conflicts of Interest and Disclosure Statement

none

## Notes

### Competing Interest Statement

The authors have declared no competing interest.

