## Supplementary material for "Basal forebrain activity predicts functional degeneration in the entorhinal cortex and decreases with Alzheimer’s Disease progression"



*****Data used in preparation of this article were obtained from the Alzheimer's Disease Neuroimaging Initiative (ADNI) database (adni.loni.usc.edu). As such, the investigators within the ADNI contributed to the design and implementation of ADNI and/or provided data but did not participate in analysis or writing of this report. A complete listing of ADNI investigators can be found at: http://adni.loni.usc.edu/wp-content/uploads/how_to_apply/ADNI_Acknowledgement_List.pdf

### Methods

#### Image acquisition

Participants were scanned at multiple sites equipped with 3-Tesla MRI scanners according to unified ADNI monitoring protocols [1]. All scanner sites had to pass strict scanner validation tests before contributing. To ensure maximum compatibility between the measurements, we adhered ADNI's recommendations including only the basic rsfMRI version of ADNI 3 since the advanced version is not compatible with ADNI-GO/2. Moreover, all participants included in our analysis were examined with the same scanner using the same type of head coil for both timepoints, T1 and T2 (https://adni.loni.usc.edu/methods/mri-tool/mri-analysis/), this led to the exclusion of 3 participants in ADNI-GO/2 and 16 in ADNI 3 data. Furthermore, for the inclusion of images, the quality criteria by ADNI were adhered by only including measurements with excellent, good, or fair quality, thus, we excluded n=8 ADNI-GO/2 and n=14 ADNI 3 data sets.

T1- weighted sagittal images were obtained using a magnetization-prepared rapid gradient echo (MP-RAGE) with ~1mm isotropic voxel size and TR(ms)/TE(ms)/flip angle(degree)=6.5-2300/2.93-3.12/9° for Philips and Siemens, 11° for GE. rsfMRI was acquired using axial echo-planar imaging (EPI). For each subject's measurement, volumes between 140 and 200 were recorded in ~3.3mm isotropic voxel resolution with a TR(ms)/TE(ms)/flip angle(degree)=3000-3025/ 30-30.001/ 80° for ADNI-GO/2, 90° for ADNI 3. Fieldmaps were included if available to correct for B0 inhomogeneities. The following protocol parameters were applied: Philips: TR(ms)/TE1(ms)/TE2(ms)/flip angle (degree)=20/2.3/4.6/10°; Siemens: TR(ms)/TE1(ms)/ TE2(ms)/flip angle (degree)=571-581/4.92/7.38/60°. For GE no fieldmapping was conducted. Detailed descriptions of image acquisition can be found on http://adni.loni.usc.edu.

#### Data preprocessing

Considering their specific scanning parameters such as TR, slice order, and volume number, all data were preprocessed with the Data Processing Assistant for Resting-State fMRI Advanced (DPARSFA, http://rfmri.org/dpabi) toolbox version 5 (release 5.2_210501), which is based on the Statistical Parametric Mapping toolbox (SPM 12, https://www.fil.ion.ucl.ac.uk/spm/) for MATLAB®. The preprocessing pipeline started with removing the first ten volumes for the signal to reach T1 equilibrium and the subject to adjust to the scanner's environment. Further steps included a) slice time correction; b) spatial realignment using a six-parameter rigid-body spatial transformation and unwarping using $B_0$ fieldmaps if available to mitigate spurious effects of head motions during the measurement; c) T1 co-registration to the mean functional image; d) CSF, gray and white matter tissue class segmentation, as well as spatial normalization using diffeomorphic anatomical registration using exponential lie algebra (DARTEL) [2] for T1 images; e) regression of nuisance variables, including CSF and white matter signal, and the Friston 24-parameter model [3] to mitigate the influence of spurious physiological effects and head motion; f) functional images were normalized to MNI space and resampled to an isotropic voxel size of 3 mm using the parameters estimated by DARTEL.

For ReHo and functional connectivity measures, the normalized images were bandpass filtered (0.01-0.1 Hz). In a next step, scrubbing was applied to detect and remove single frames affected by head motion, so-called "bad" time points. Frames exceeding a frame-to-frame displacement of 1 mm have been removed along with the previous and the two subsequent frames [4]. Only scans with >70% remaining EPI volumes after scrubbing were included to ensure a sufficiently high number of volumes for the subsequent analysis [5]. Consequently, we excluded these participants who did not meet this criterion (ADNI-GO/2: n=16, ADNI 3: n=4). For fALFF, no filtering and scrubbing were applied [6].

To reduce the influence of excessive head motion, participants exhibiting more than 3.0 mm of maximum movement and a 3.0-degree rotation angle were discarded (n=3 for ADNI 3).

Further, Images were visually inspected after co-registration, segmentation, and normalization to guarantee high quality. This included a specific focus on signal loss and artifacts in the regions of interest (NbM, EC) by overlaying the ROI mask in standardized space; especially, the EC represents a region that might often be affected by artifacts [7]. This check resulted in a final sample size of n=71 (Table 1).

Since head motion strongly influences functional connectivity measures [4], we compared head motion in a dependent t-test with CSF group as a grouping factor to ensure no confounding effect on groups. Consequently, the comparison in the 6 parameters of rigid body transformation of maximum and mean motion revealed no significant differences between normal CSF and abnormal CSF, except for measurement 2 the mean rotation in z-axis ($t(69)=2.425$, $p=0.018$). Furthermore, the root mean square of head motion, same as the relative root mean square of framewise displacement (FD) Vanijk showed no significant differences between the two groups. In measurement 1 there was a significant difference between nCSF and aCSF in the mean FD Power ($t(69)=-2.276$, $p=0.026$) and in measurement 2 ($t(69)=-3.228$, $p=0.02$). Because the groups only differed regarding their mean FD Power and the other tests revealed no significant differences, we minimized the influence of head motions by excluding participants with excessive head motions and utilizing scrubbing.

#### Region of interest definition

The regions of interest NbM and EC were created in MNI space combined across both hemispheres with the SPM Anatomy Toolbox Version 3.0 [8–10] (available from https://www.fz-juelich.de/en/inm/inm-7/resources/jubrain-anatomy-toolbox, Fig. 1A) The NbM is defined as the Ch4 according to previous published probabilistic maps [11]. The EC was defined based on the previously published probability map [12] .

We used the toolbox MarsBaR [13] to extract the mean rsfMRI signal intensity for each ROI at a threshold of 50% probability and each subject individually.

#### rsfMRI analyses

**The fractional amplitude of low-frequency fluctuations analysis (fALFF)**

Spontaneous local brain activity based on the amplitude of BOLD signals can be assessed by the amplitude of low-frequency fluctuations (ALFF) and its improved measure of fractional amplitude of low-frequency fluctuations (fALFF) [14–16]. ALFF assesses the amplitude in the low-frequency range, and fALFF represents the ratio of total amplitude within the low-frequency range (here, 0.01-0.1 Hz) to the amplitude across the entire detectable frequency range. Importantly, fALFF shows a higher specificity with regard to the detection of local spontaneous brain activity and is more robust against nonspecific signals (e.g., physiological noise) as compared to ALFF [15,16]. Therefore, fALFF is recommended [17] and used here.

**Regional Homogeneity (ReHo)**

Regional homogeneity (ReHo) refers to local connectivity within brain regions to measure regional synchronization among brain voxels and their neighboring voxels [18]. The central assumption of ReHo is that structural neighboring voxels represent a functional homogeneity of time series defining it a network centrality metric [19]. In this study, ReHo is quantified by Kendall's coefficient of concordance (KCC) [20] of a given voxel with its nearest neighbors. Thus, a larger ReHo value indicates higher local synchronization.

**ROI to ROI functional connectivity**

Functional connectivity is defined by a temporal dependence across anatomically separated brain regions regarding patterns of neuronal activity [14,21,22]. Functional connectivity is supposed to reflect functional communication and a shared function between brain regions [23]. Therefore, Pearson's correlation coefficient between both time courses extracted from the NbM and EC was calculated and a Fisher's z transformation was applied.

### Results

#### ReHo

**Regions differ regarding their regional homogeneity (ReHo), however, not CSF group**

The 2x2 mixed ANCOVA revealed a significant main effect of region ($F(1,69)=8.387$, $p=0.005$, partial $\eta^2=0.108$, Fig. S1A), no main effect of group ($F(1,63)=0.003$, $p=0.960$, partial $\eta^2=0.000$, Fig. S1A) and no significant interaction between group and region ($F(1,63)=1.706$, $p=0.196$ , partial $\eta^2=0.026$, Fig. S1A).

**Annual percentage signal change in ReHo does not differentiate between CSF groups or regions**

We used a 2x2 mixed ANCOVA to investigate whether the longitudinal indices of APSC in ReHo of the NbM and EC differentiated between CSF normal vs. abnormal groups. There was no significant main effect of region ($F(1,69)=0.001$, $p=0.974$, partial $\eta^2=0.000$, Fig. S1B), no main effect of CSF group ($F(1,63)=0.218$, $p=0.642$, partial $\eta^2=0.003$, Fig. S1B), and no interaction of region and CSF group ($F(1,63)=0.394$, $p=0.532$ , partial $\eta^2=0.006$, Fig. S1B).

**The baseline signal in NbM does not predict the annual percentage signal change in ReHo of EC**

The robust regression modeling did not reveal any significant results (see Fig. S2, Table S2).

#### CSF group does not moderate the functional spread of degeneration in ReHo

The moderation analysis included baseline ReHo NbM as independent variable, ReHo EC APSC as dependent variable and CSF group as moderator. The model was not statistically significant ($R^2$=0.2135, F(9,61)=1.2677, p=0.2728), with no significant direct effect of NbM→EC (t(61)=-0.5922, p=0.5559), and no significant moderator effect (t(61)=-0.5067, p=0.6142), which is in line with the robust regression analysis.

The moderation analysis included baseline ReHo EC as independent variable, ReHo NbM APSC as dependent variable and CSF group as moderator. The model was also not statistically significant ($R^2$=0.0544, F(9,61)=0.3712, p=0.9445), with no significant direct effect of EC→NbM (t(61)=-0.4040, p=0.6876), and no significant moderator effect (t(61)=-0.7685, p=0.4451, which, again, is in line with the robust regression analysis.

#### Functional connectivity between NbM and EC

**NbM and EC show functional connectivity independent from CSF status**

To investigate functional connectivity between NbM and EC, we used a 2x2 mixed ANCOVA with time (timepoint 1 and timepoint 2) and group (aCSF, nCSF) as within and between group variables. There was no significant effect of time (F(1,69)=1.221, p=0.273 , partial $\eta^2$=0.017, Fig. S1C), no main effect of CSF group (F(1,63)=1.467, p=0.230, partial $\eta^2$=0.023, Fig. S1C), and no significant interaction between time and group (F(1,63)=0.571, p=0.453 , partial $\eta^2$=0.009, Fig. S1C).

However, given our a priori hypotheses of a functional connectivity between both regions, we carried out t-tests for each time point separately across both groups. The revealed a significant effect, and therefore functional connectivity, for timepoint 1 in nCSF (t(36)=2.667, p=0.011) and timepoint 2 (t(36)=3.054, p=0.004). In aCSF there was only a borderline significant effect at timepoint 1 ($t(33)=1.999$, $p=0.054$) but a highly significant effect in timepoint 2 ($t(33)=3.535$, $p=0.001$).

#### Supplementary material figure S1

ReHo signals and annual percentage change and functional connectivity between NbM and EC

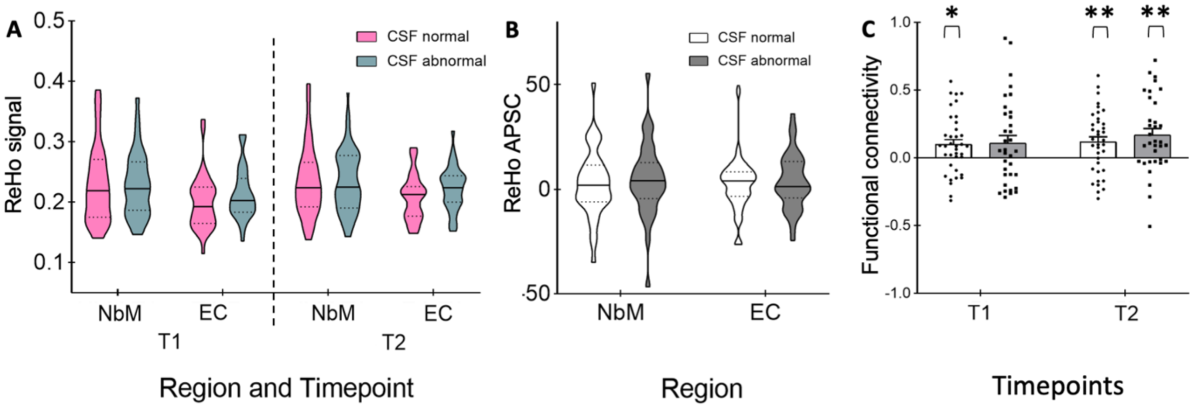


**Figure S1.** A) Violin plots representing the baseline signals (T1) and the signals at the follow-up measurement (T2) in ReHo and B) showing the annual percentage signal change (APSC) in both regions in ReHo. The horizontal lines show the median and the dotted lines the interquartile range.

The functional connectivity between NbM and EC at T1 and T2 represented in bar plots in C). Here the means are shown with standard error bars. *p<0.05, **p<0.01.

**Supplementary material figure S2**

Robust regression models in ReHo

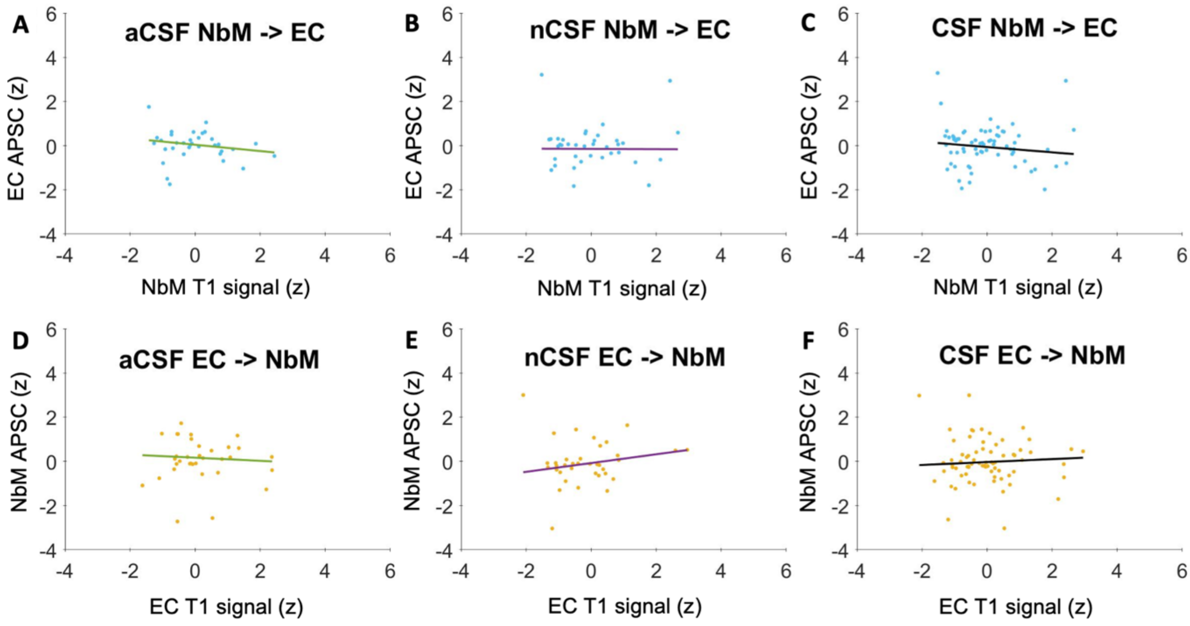


**Figure S2.** Adjusted for covariates response plots for the robust regression models in ReHo for A) – C) predictive spread in NbM→EC and D) – F) predictive spread in EC→NbM in aCSF (A) and D)) and nCSF (B) and E)) as well as CSF included as a categorical predictor (C) and F)). The x-axis shows the region's z-scored baseline signal, whereas the y-axis reflects the z-scored annual percentage signal change (APSC) in ReHo.

**Supplementary material table S1**

The output of the robust regression models of fALFF with the covariates age, sex, ADNI cohort, scanner and education.

**aCSF NbM → EC (n= 34)**

| | Estimate | SE | t | p |
|---|---|---|---|---|
| Intercept | 3.74 | 2.88 | 1.3 | 0.205 |
| Baseline NbM | -0.3 | 0.22 | -1.36 | 0.185 |
| Sex | -0.05 | 0.45 | -0.11 | 0.91 |
| Age | -0.06 | 0.03 | -2.28 | 0.031 |
| ADNI Group | 1.28 | 1.1 | 1.17 | 0.251 |
| Scanner: Siemens | -0.63 | 1.24 | -0.51 | 0.615 |
| Scanner: GE | -1.8 | 1.23 | -1.46 | 0.155 |
| Education | 0.06 | 0.1 | 0.6 | 0.551 |

R-Squared: 0.263; Adjusted R- Squared: 0.0649; df= 26; F-statistics vs. constant model: 1.33; p-value: 0.277

**nCSF NbM → EC (n= 37)**

| | Estimate | SE | t | p |
|---|---|---|---|---|
| Intercept | 0.95 | 2.15 | 0.44 | 0.662 |
| Baseline NbM | -0.43 | 0.15 | -2.81 | 0.009 |
| Sex | -0.11 | 0.33 | -0.34 | 0.738 |
| Age | -0.01 | 0.02 | -0.38 | 0.708 |
| ADNI Group | 0.13 | 0.46 | 0.28 | 0.78 |
| Scanner: Siemens | 0.42 | 0.48 | 0.88 | 0.388 |

| | | | | |
|---|---|---|---|---|
| Scanner: GE | -0.34 | 0.54 | -0.63 | 0.536 |
| Education | -0.02 | 0.07 | -0.24 | 0.813 |

R-Squared: 0.296; Adjusted R- Squared: 0.126; df= 29; F-statistics vs. constant model: 1.74; p-value: 0.138

**aCSF EC → NbM (n= 34)**

| | Estimate | SE | t | p |
|---|---|---|---|---|
| Intercept | -2.24 | 3.11 | -0.72 | 0.478 |
| Baseline EC | -0.26 | 0.25 | -1.03 | 0.314 |
| Sex | 0.06 | 0.53 | 0.12 | 0.908 |
| Age | 0.02 | 0.03 | 0.5 | 0.623 |
| ADNI Group | 1.28 | 1.2 | 1.07 | 0.296 |
| Scanner: Siemens | -0.83 | 1.36 | -0.61 | 0.545 |
| Scanner: GE | -1.08 | 1.36 | -0.8 | 0.433 |
| Education | 0.05 | 0.11 | 0.51 | 0.614 |

R-Squared: 0.137; Adjusted R- Squared: -0.0929; df= 26; F-statistics vs. constant model: 0.587; p-value: 0.76

**nCSF EC → NbM (n= 37)**

| | Estimate | SE | t | p |
|---|---|---|---|---|
| Intercept | 2.15 | 1.89 | 1.14 | 0.264 |
| Baseline EC | -0.21 | 0.13 | -1.58 | 0.125 |
| Sex | 0.03 | 0.3 | 0.09 | 0.926 |
| Age | -0.02 | 0.02 | -1.14 | 0.262 |
| ADNI Group | -0.01 | 0.4 | -0.02 | 0.984 |
| Scanner: Siemens | 0.54 | 0.42 | 1.27 | 0.214 |
| Scanner: GE | -0.03 | 0.47 | -0.07 | 0.944 |
| Education | -0.04 | 0.06 | -0.58 | 0.565 |

R-Squared: 0.175; Adjusted R- Squared: -0.0239; df= 29; F-statistics vs. constant model: 0.88; p-value: 0.534

208

**With CSF NbM → EC (n= 71)**

| | Estimate | SE | t | p |
|---|---|---|---|---|
| Intercept | 2.2 | 1.61 | 1.36 | 0.178 |
| Baseline NbM | -0.38 | 0.12 | -3.14 | 0.003 |
| Sex | -0.19 | 0.25 | -0.78 | 0.44 |
| CSF | 0.18 | 0.26 | 0.72 | 0.477 |
| Age | -0.03 | 0.02 | -1.99 | 0.051 |
| ADNI Group | 0.32 | 0.39 | 0.81 | 0.42 |
| Scanner: Siemens | 0.12 | 0.43 | 0.27 | 0.788 |
| Scanner: GE | -0.69 | 0.47 | -1.48 | 0.145 |

| | | | | |
|---|---|---|---|---|
| Education | 0.01 | 0.05 | 0.28 | 0.783 |

R-Squared: 0.235; Adjusted R- Squared: 0.137; df= 62; F-statistics vs. constant model: 2.39; p-value: 0.026

**With CSF EC → NbM (n= 71)**

| | Estimate | SE | t | p |
|---|---|---|---|---|
| Intercept | 0.04 | 1.62 | 0.03 | 0.98 |
| Baseline EC | -0.21 | 0.11 | -1.8 | 0.077 |
| Sex | -0.04 | 0.25 | -0.17 | 0.866 |
| CSF | 0.08 | 0.25 | 0.33 | 0.741 |
| Age | -0.004 | 0.02 | -0.22 | 0.823 |
| ADNI Group | 0.16 | 0.38 | 0.41 | 0.685 |
| Scanner: Siemens | 0.32 | 0.43 | 0.73 | 0.466 |
| Scanner: GE | -0.06 | 0.47 | -0.12 | 0.905 |
| Education | 0.003 | 0.05 | 0.06 | 0.951 |

R-Squared: 0.0884; Adjusted R- Squared: -0.0293; df= 62; F-statistics vs. constant model: 0.751; p-value: 0.646

**Supplementary material table S2**

The output of the robust regression models of ReHo with the covariates age, sex, ADNI cohort, scanner and education.

**aCSF NbM → EC (n= 34)**

| | Estimate | SE | t | p |
|---|---|---|---|---|
| Intercept | -5.07 | 1.97 | -2.58 | 0.016 |
| Baseline NbM | -0.14 | 0.15 | -0.98 | 0.337 |
| Sex | 0.4 | 0.3 | 1.31 | 0.201 |
| Age | 0.04 | 0.02 | 2.28 | 0.031 |
| ADNI Group | -1.33 | 0.75 | -1.78 | 0.087 |
| Scanner: Siemens | 1.12 | 0.85 | 1.32 | 0.198 |
| Scanner: GE | 3.06 | 0.84 | 3.66 | 0.001 |
| Education | 0.1 | 0.07 | 1.53 | 0.137 |

R-Squared: 0.525; Adjusted R- Squared: 0.397; df= 26; F-statistics vs. constant model: 4.1; p-value: 0.00373

**nCSF NbM→ EC (n= 37)**

| | Estimate | SE | t | p |
|---|---|---|---|---|
| Intercept | 0.83 | 2.3 | 0.36 | 0.722 |
| Baseline NbM | -0.01 | 0.15 | -0.04 | 0.971 |
| Sex | 0.12 | 0.36 | 0.33 | 0.743 |
| Age | -0.03 | 0.03 | -1.03 | 0.313 |
| ADNI Group | -0.09 | 0.47 | -0.19 | 0.851 |

| | | | | |
|---|---|---|---|---|
| Scanner: Siemens | -0.43 | 0.51 | -0.86 | 0.399 |
| Scanner: GE | 0.34 | 0.59 | 0.57 | 0.57 |
| Education | 0.06 | 0.07 | 0.78 | 0.442 |

R-Squared: 0.145; Adjusted R- Squared: -0.0608; df= 29; F-statistics vs. constant model: 0.705; p-value: 0.668

**aCSF EC → NbM (n= 34)**

| | Estimate | SE | t | p |
|---|---|---|---|---|
| Intercept | -2.38 | 3.15 | -0.76 | 0.455 |
| Baseline EC | -0.07 | 0.21 | -0.33 | 0.744 |
| Sex | 0.14 | 0.46 | 0.31 | 0.757 |
| Age | 0.02 | 0.03 | 0.64 | 0.526 |
| ADNI Group | 0.08 | 1.11 | 0.07 | 0.946 |
| Scanner: Siemens | 1.87 | 1.27 | 1.48 | 0.152 |
| Scanner: GE | 0.49 | 1.25 | 0.39 | 0.7 |
| Education | 0.05 | 0.1 | 0.52 | 0.608 |

R-Squared: 0.316; Adjusted R- Squared: 0.131; df= 26; F-statistics vs. constant model: 1.71; p-value: 0.15

**nCSF EC → NbM (n= 37)**

| | Estimate | SE | t | p |
|---|---|---|---|---|
| Intercept | 1.65 | 2.4 | 0.69 | 0.497 |
| Baseline EC | 0.2 | 0.16 | 1.21 | 0.237 |
| Sex | -0.06 | 0.37 | -0.17 | 0.863 |
| Age | -0.02 | 0.03 | -0.9 | 0.377 |
| ADNI Group | 0.56 | 0.48 | 1.16 | 0.254 |
| Scanner: Siemens | -0.36 | 0.52 | -0.68 | 0.501 |
| Scanner: GE | 0.15 | 0.59 | 0.25 | 0.804 |
| Education | -0.01 | 0.08 | -0.17 | 0.863 |

R-Squared: 0.139; Adjusted R- Squared: -0.0688; df= 29; F-statistics vs. constant model: 0.669; p-value: 0.696

**With CSF NbM → EC (n= 71)**

| | Estimate | SE | t | p |
|---|---|---|---|---|
| Intercept | -2.02 | 1.6 | -1.27 | 0.21 |
| Baseline NbM | -0.12 | 0.11 | -1.05 | 0.297 |
| Sex | 0.19 | 0.24 | 0.78 | 0.441 |
| CSF | 0.1 | 0.24 | 0.42 | 0.678 |
| Age | 0.02 | 0.02 | 0.89 | 0.376 |
| ADNI Group | -0.22 | 0.38 | -0.58 | 0.566 |
| Scanner: Siemens | -0.13 | 0.42 | -0.3 | 0.763 |
| Scanner: GE | 1.18 | 0.47 | 2.525 | 0.014 |

| | | | | |
|---|---|---|---|---|
| Education | 0.04 | 0.05 | 0.84 | 0.403 |

R-Squared: 0.169; Adjusted R- Squared: 0.0623; df= 62; F-statistics vs. constant model: 1.58; p-value: 0.149

**With CSF EC→ NbM (n= 71)**

| | Estimate | SE | t | p |
|---|---|---|---|---|
| Intercept | 0.14 | 1.77 | 0.08 | 0.939 |
| Baseline EC | 0.07 | 0.12 | 0.54 | 0.59 |
| Sex | -0.1 | 0.27 | -0.36 | 0.723 |
| CSF | 0.26 | 0.26 | 1.01 | 0.317 |
| Age | -0.01 | 0.02 | -0.3 | 0.768 |
| ADNI Group | 0.41 | 0.42 | 1.0 | 0.323 |
| Scanner: Siemens | -0.29 | 0.47 | -0.63 | 0.531 |
| Scanner: GE | 0.17 | 0.51 | 0.33 | 0.743 |
| Education | 0.002 | 0.06 | 0.04 | 0.972 |

R-Squared: 0.055; Adjusted R- Squared: -0.669; df= 62; F-statistics vs. constant model: 0.451; p-value: 0.885
